# Genomics 2 Proteins portal: A resource and discovery tool for linking genetic screening outputs to protein sequences and structures

**DOI:** 10.1101/2024.01.02.573913

**Authors:** Seulki Kwon, Jordan Safer, Duyen T. Nguyen, David Hoksza, Patrick May, Jeremy A. Arbesfeld, Alan F. Rubin, Arthur J Campbell, Alex Burgin, Sumaiya Iqbal

**Affiliations:** Broad Institute of MIT and Harvard, Center for the Development of Therapeutics, Cambridge, MA 02142; Broad Institute of MIT and Harvard, PATTERN, Cambridge, MA 02142; Department of Software Engineering, Faculty of Mathematics and Physics, Charles University, Prague, Czech Republic; Luxembourg Centre for Systems Biomedicine, University of Luxembourg, Esch-sur-Alzette, Luxembourg; The Steve and Cindy Rasmussen Institute for Genomic Medicine, Nationwide Children’s Hospital, Columbus, OH; Walter and Eliza Hall Institute of Medical Research, Parkville, Victoria, Australia; Department of Medical Biology, University of Melbourne, Melbourne, Victoria, Australia; Program in Medical and Population Genetics, Broad Institute of MIT and Harvard, Cambridge, MA 02142; Analytic and Translational Genetics Unit, Massachusetts General Hospital, Boston, MA 02114; Cancer Data Sciences, Dana-Farber/Harvard Cancer Center, Boston, MA 02215

## Abstract

Recent advances in AI-based methods have revolutionized the field of structural biology. Concomitantly, high-throughput sequencing and functional genomics technologies have enabled the detection and generation of variants at an unprecedented scale. However, efficient tools and resources are needed to link these two disparate data types – to “map” variants onto protein structures, to better understand how the variation causes disease and thereby design therapeutics. Here we present the Genomics 2 Proteins Portal (G2P; g2p.broadinstitute.org/): a human proteome-wide resource that maps 19,996,443 genetic variants onto 42,413 protein sequences and 77,923 structures, with a comprehensive set of structural and functional features. Additionally, the G2P portal generalizes the capability of linking genomics to proteins beyond databases by allowing users to interactively upload protein residue-wise annotations (variants, scores, etc.) as well as the protein structure to establish the connection. The portal serves as an easy-to-use discovery tool for researchers and scientists to hypothesize the structure-function relationship between natural or synthetic variations and their molecular phenotype.

---

We live in the era of big biological data where deep learning methods for protein structure prediction have opened the door for a data-driven revolution in structural biology, making millions of high-quality predicted protein structures accessible to the biomedical community^1–7^. At the same time, technological advancements in Cryo-electron microscopy (Cryo-EM) and other experimental methods are leading to a burst of high-resolution protein structures and macromolecular assemblies^8–11^. These advances come at a time when an unprecedented number of genetic variants in the general population and those associated with monogenic and complex diseases have been identified and accumulated in multiple databases^12–17^. Concomitantly, advances in functional genomics approaches (e.g., base-editing^18,19^, prime-editing^20^, Perturb-Seq^21^) have enabled the generation of synthetic mutations and the quantification of their functional impact in different *in vitro* and *in vivo* models. Investigating a target gene by mapping natural or synthetic variants in the context of protein structure provides valuable molecular-level insights and helps hypothesize the structure-function mechanism of the variant.

Challenges remain, however, in connecting genomic data to protein structural data, due to the complexity introduced by diverse RNA transcripts and protein isoforms originating from a single DNA sequence^22,23^. Determining the relevant transcript and protein isoforms for a given variant relies on precise transcript-isoform mapping, which is crucial for assessing variant impacts on proteins correctly^24^. Moreover, there is a technical hurdle in reconciling disparate data formats between genomic annotations of variants such as rsIDs^25^ or HGVS notations^26^, and those of protein data, which are featured on amino acid sequences and their spatial coordinates^27,28^ in structures, often only available for fragments of the full-length protein. This discrepancy in naming and formatting requires interdisciplinary knowledge or collaborations between researchers from different disciplines, including genetics, structural biology, and computational biology, to align and harmonize the data. Therefore, the linkage of human proteome-wide genetic and structural data at scale necessitates efficient computational methods capable of accurately matching variants to their corresponding protein structures to better understand how the variation causes disease and thereby design better therapeutics, which is precisely why we developed the Genomics 2 Proteins portal (G2P; g2p.broadinstitute.org).

The G2P portal is an integrated bioinformatic tool to dynamically query, retrieve, and connect genetic variants and transcripts to protein sequence annotations and structures, wrapped within an interactive web interface with visualization functions. By programmatic access and linking of APIs, over 20 million genetic variants within all human protein-coding genes from public databases (ClinVar^12^, HGMD^13^; Human Gene Mutation Database, and gnomAD^15^; Genome Aggregation Database) are aggregated and annotated within protein sequences and structures with comprehensive protein feature reports. By exploiting experimentally solved (PDB^8^; Protein Data Bank) and predicted (AlphaFold database^29^; AlphaFoldDB) protein structures, G2P covers 99.2% (20,132 out of 20,292) of all human proteins corresponding to 77,923 structures.

## Results

### The Genomics 2 Proteins (G2P) Bioinformatic Framework

The G2P portal is constructed upon an integrated bioinformatic method for linking genetic variants from genes and transcripts to protein sequences and structures across the entire human proteome. We integrated public databases to build a new API for seamless mapping of identifiers for genes (HGNC^30^; HUGO Gene Nomenclature Committee), transcripts (Ensembl^31^; the Ensembl genome browser and RefSeq^32^; NCBI Reference Sequence Database), protein sequences (UniProtKB^28^; UniProt KnowledgeBase), and structures (PDB^8^ and AlphaFoldDB^29^) (API: **G**enomics **2 P**roteins **3D** or G2P3D, see **Methods**: Construction of G2P3D API). The G2P3D API currently links 20,292 human genes^30,33^ that encode 20,242 UniProtKB^28^ protein identifiers (UniProtAC) corresponding to 42,413 protein isoforms, via 53,607 Ensembl^31^ transcripts and 57,543 RefSeq^32^ transcripts, to 77,923 protein structures (58,027 from the PDB^8^ and 19,896 from AlphaFoldDB^29^). A schematic of the portal and data flow via the G2P3D API is illustrated in **Fig. 1**.

**Fig. 1.**
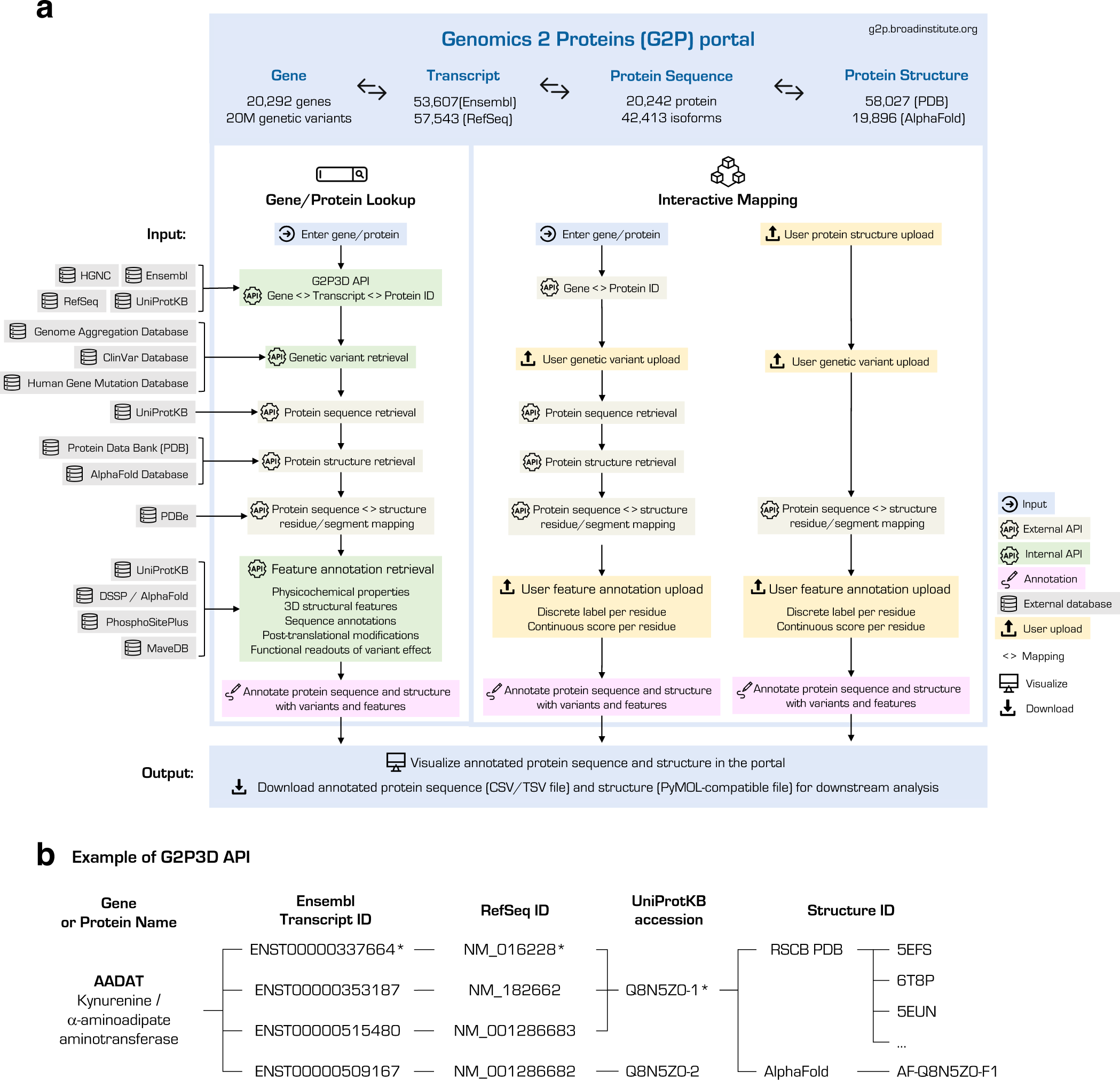
The Bioinformatic Framework of the Genomics 2 Protein (G2P) portal. **(a)** The schematic overview of data and method integration in the G2P portal and its two main modules: Gene/Protein lookup and Interactive mapping. In the Gene/Protein lookup module, the connections across identifiers of human genes, transcripts, protein sequences and structures were established using an in-house API: G2P3D, for the entire human proteome (see **Methods** for details). Variants from databases, such as gnomAD^15^, ClinVar^12^, and HGMD^13^, were subsequently mapped onto protein sequences and structures upon dynamically querying UniProtKB and structure databases (PDB^8^ and AlphaFoldDB^29^), respectively. Additionally, protein feature annotations were fetched and calculated from various databases (UniProtKB^28^, DSSP^43^, PhosphoSitePlus^34^, etc.). All annotated protein sequences and structures with variants and features are viewable on the portal and downloadable in interoperable formats for further analyses. In the Interactive mapping module of the portal, users can upload protein residue-wise annotations of variants and additional features and perform linking genetic data to protein structural data. Users can access this module by starting from a gene and by uploading an in-house protein structure. **(b)** An example of G2P3D API output; the API links human genes (HGNC^30^) to transcripts (Ensembl^31^ and RefSeq^32^) to protein sequences (UniProtKB) and structures (PDB and AlphaFoldDB). In this example, *AADAT* has 4 Ensembl transcripts and 4 RefSeq transcripts; 3 pairs of Ensembl-RefSeq transcripts encode the canonical protein isoform (Q8N5Z0-1*) and the remaining one transcript (ENST00000509167/NM_001286682) corresponds to non-canonical protein isoform, Q8N5Z0-2. The canonical protein isoform is further dynamically linked to multiple available PDB structures and the AlphaFold structure. In the portal, variants are mapped onto both canonical and non-canonical protein isoforms. Only canonical protein isoform variants are mapped to available protein structures.

Multi-omics data aggregated in the portal showed that about 47% of all human genes (9,671 out of 20,292) have one unique isoform (e.g., Gene/UniProtAC: *A2M*/P01023), and the other 10,621 have on average 3.1 isoforms (ranging from 2 to 37) by alternative splicing (e.g., *CACNA1C*/Q13936-1;…;Q13936-37). Furthermore, 75.7% of all protein isoforms (25,523 out of 33,720) have a single Ensembl transcript, whereas the remaining 24.3% (8,197 isoforms) have, on average, 3.0 Ensembl transcript. These statistics vary for RefSeq transcript database, with 62.5% of protein isoforms having a single RefSeq transcript, whereas 9,392 isoforms (27.9%) have, on average, 3.7 RefSeq transcripts and the rest of the 3,243 isoforms (9.6%) having no RefSeq transcript.

We noted 53 genes (out of 20,292) with multiple UniProtKB identifiers, ranging from two to five UniProtACs (e.g. *AKAP7/O43687;Q9P0M2).* At the same time, 73 UniProtACs (out of 20,242) correspond to multiple genes (from 2 to 14), e.g., *CKMT1A;CKMT1B*/P12532. For the former case, users can select a UniProtKB identifier in the G2P portal for the gene of interest to explore variants on the selected protein’s sequence and structure. For 17,679 genes (out of 20,292, 87.1%), canonical transcript translates into canonical protein isoforms (e.g. *AAR2* or *ZYX*). For 1,836 genes (9.0%), canonical transcript and canonical protein isoform do not pair with each other (e.g. *A1CF* or *AAR2*). For 774 genes (3.8%), no protein-coding transcript is available (e.g. *ABCA13* or *ABI2*). Regarding protein structural data, 7,977 out of 20,292 genes (39.3%) have experimentally solved 3D structures of the encoded proteins in PDB. In contrast, 20,027 genes (98.6%) have a predicted structure in AlphaFoldDB. In total, 20,132 genes (99.2%) have either PDB or AlphaFold structures (77,923 structures corresponding to 20004 proteins).

We implemented the above-mentioned bioinformatic framework in two distinct modules of the G2P portal (**Fig. 1a**): (1) Gene/Protein Lookup: a human proteome-wide resource for users to link genetic variants from transcripts to protein sequences and structures and (2) Interactive Mapping: a tool for users to analyze their own data, which is thereby not limited to publicly available variants or protein structures. In the Gene/Protein Lookup module, ∼20M variants aggregated from different databases were linked to their corresponding UniProtAC identifiers using the in-house G2P3D API (**Fig. 1b**) and then mapped onto its amino acid position upon dynamic retrieval of the protein sequence and structure (see **Methods:** Variant and feature mapping onto proteins). In addition to variants, a wide set of protein residue-wise annotations (referred to as protein features) are computed and aggregated in the portal, such as protein sequence annotation from UniProtKB, structural features, post-translational modification (PTM) information from PhosphoSitePlus^34^, readouts of multiplexed assays of variants effect from MaveDB^35^, etc. (**Fig. 1a**). Finally, variants are mapped onto protein sequence and structure simultaneously with protein features, as it is often useful to find out the loss of important features due to mutation for informed variant prioritization.

### Resources in the G2P portal

The Gene/Protein Lookup module of the G2P portal contains variants, protein structures and protein features for 20,292 human genes and encoded proteins (**Supplementary Table 1).** Genes and proteins are classified by HGNC^30^ gene family and PANTHER^36^ protein class (**Methods**: Gene family and protein class annotations in the G2P portal; **Supplementary Fig. 1**). About 75% of genes were annotated to be a part of an HGNC gene family (834 families; **Supplementary Table 2**). Additionally, for about 69% of proteins, the protein functional class annotations were retrievable from the PANTHER knowledgebase^36^ (24 classes; **Supplementary Table 3).**

#### Variant data

We applied the G2P bioinformatic method to variants in three human genetic variation databases: the genome aggregation database^15^ (gnomAD), Clinically identified variants^12^ (ClinVar), and the Human Gene Mutation Database^13^ (HGMD). Throughout the paper, we refer to variants from these databases as gnomAD, ClinVar, and HGMD variants. A total of 20 million variants leading to protein consequences, covering the entire protein-coding genome, were aggregated (see **Methods**: Variant aggregation) and mapped to their corresponding protein sequence and structure positions. Variants were grouped based on their protein consequence: missense, nonsense, synonymous, frameshift, in-frame indel, and others, and visualized as separate tracks in the protein sequence viewer of the portal, with a rich set of protein structural and functional features (described below).

From gnomAD (v2.1.1), we obtained 18,014,632 protein-coding variants annotated in 41,570 transcripts corresponding to 18,723 human genes. About 63% of all protein-coding variants in gnomAD were missense, i.e., a single nucleotide variation (SNV) leading to a single amino acid substitution (**Fig. 2a**). Additionally, 31% and 2% of the variants were synonymous (no change in the protein) and nonsense (resulting in truncation of the protein), respectively. About 4% of gnomAD variants are non-SNV leading to frameshifts, in-frame insertions/deletions, etc. We incorporated allele frequency (AF) and allele count (AC) filters in the portal for users to apply for customized mapping of variants of a certain frequency. Most gnomAD variants (> ∼97%) are very rare (**Fig. 2b**), and the fraction of common variants is the largest for synonymous variants (0.78%).

**Fig. 2.**
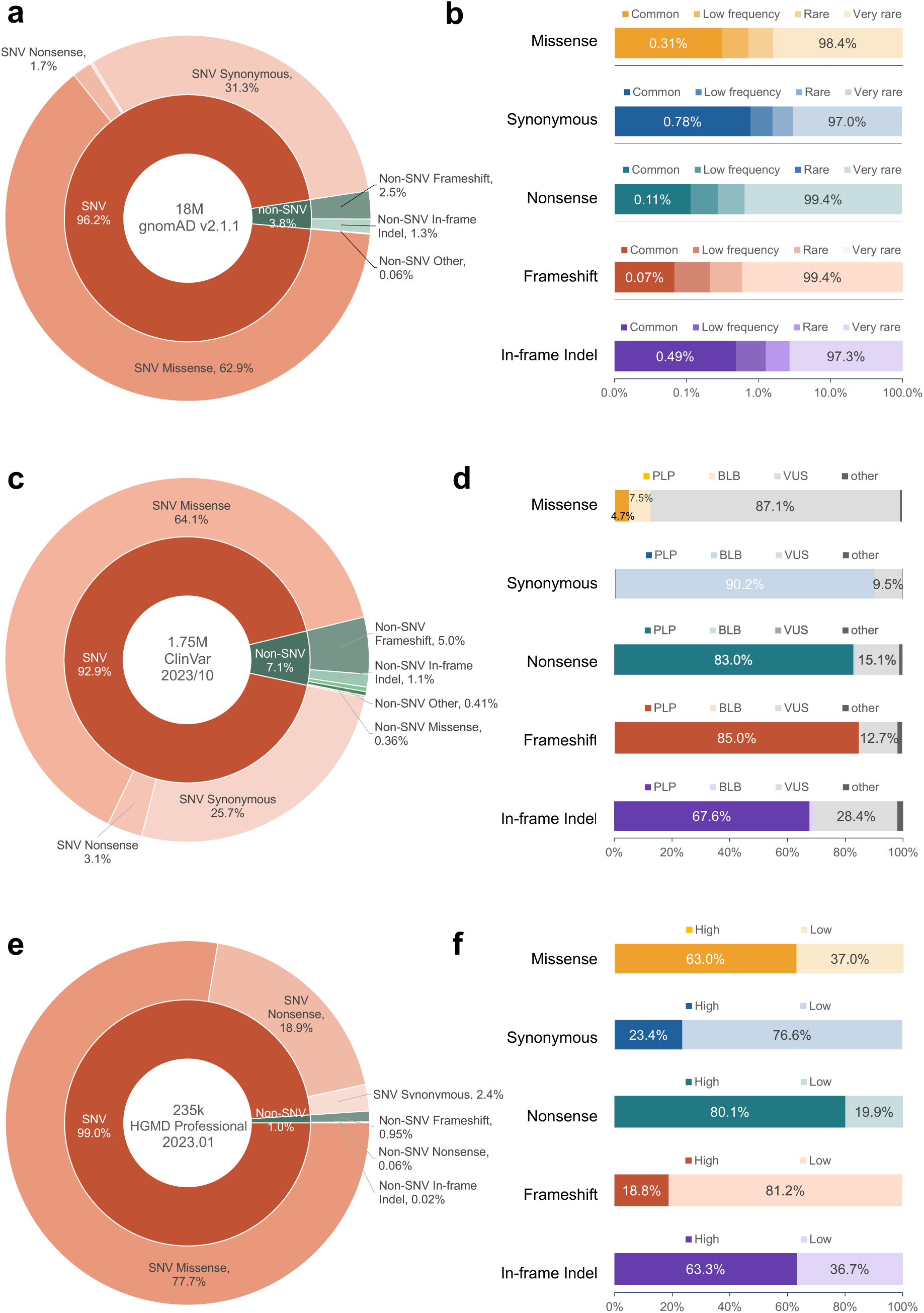
Statistics of variants from gnomAD, ClinVar, and HGMD databases aggregated in the G2P portal. **(a)** Distribution of variant types (SNV; single nucleotide variation versus non-SNV; insertion, deletion, inversion, etc.) and associated protein consequences (missense, synonymous, nonsense, frameshift, in-frame indel, and others) among 18 million protein-coding variants in gnomAD. **(b)** Fraction of gnomAD variants categorized by allele frequencies: *Very rare*; AF < 0.1%, *Rare*; 0.1% ≤ AF < 0.5%, *Low frequency*; 0.5% ≤ AF < 5%, and *Common*; AF ≥ 5%. The distributions of each allele frequency group are illustrated across different protein consequences (missense, synonymous, nonsense, frameshift, and in-frame indel). **(c)** Distribution of variant types and associated protein consequences among 1.7 million protein-coding variants in ClinVar. **(d)** Distributions of the clinical significance of ClinVar variants (PLP; pathogenic/likely pathogenic, BLB; benign/likely benign, VUS/CI; uncertain significance and conflicting interpretation, and others) displayed across different protein consequences. **(e)** Distribution of variant types and associated protein consequences among 235k HGMD variants. **(f)** Distribution of confidence levels (*High* or *Low*) for HGMD variants across different protein consequences. The scale of the x-axes for (b), (d), and (f) is linear.

It is often useful to map population (or putatively benign) variants and disease variants on proteins simultaneously to explore their relative local enrichment or depletion^37–41^. Therefore, alongside the gnomAD variants, we aggregated 1,749,628 protein-coding variants in 18,180 genes from ClinVar (10/2023 release), and 232,183 disease-causing mutations in 12,522 genes from HGMD (professional 2023.01). ClinVar variant mapping can be filtered based on the clinical significance (**Fig. 2c-d**); In the G2P portal and throughout this paper, we discussed four types of clinical significance: benign/likely-benign (BLB), pathogenic/likely-pathogenic (PLP), variants of uncertain significance and conflicting interpretation (VUS/CI), and others. **Fig. 2d** illustrates the distribution of the clinical significance of ClinVar variants across different protein consequences. As expected, the proportion of BLB and PLP variants are inversely correlated across synonymous (90.2% BLB and 0.13% PLP) and nonsense mutations (0.79% BLB and 83.0% PLP), whereas missense mutations hold the highest fraction of VUS/CI (87.1%). Of HGMD variants, 99% lead to missense/nonsense mutations (**Fig. 2e**). HGMD provides a confidence ascertainment (*High* or *Low*) of the disease mutations; synonymous mutations hold a larger fraction of low-confidence mutations (76.6%) and nonsense mutations hold the largest fraction of high-confidence mutations (80.1%; **Fig. 2f**).

Variants annotated are mapped to canonical and non-canonical protein isoforms in the portal. About 53%, 86%, 89% of gnomAD, ClinVar, and HGMD variants were annotated on the canonical protein isoforms as defined in UniProtKB (**Supplementary Fig. 2**). We found the distribution of database-specific groups, i.e., allele frequency of gnomAD variants, clinical significance of ClinVar variants, and confidence of HGMD variants, were largely unchanged across canonical versus non-canonical protein isoforms. The grouping of variants also enabled us to observe the distribution of variant types across different genes and protein classes (**Supplementary Fig. 3**).

#### Protein structural data

Human proteome-wide genetic variants aggregated in the G2P portal are mapped onto three-dimensional (3D) structures of canonical protein isoforms available in the PDB and AlphaFoldDB. G2P dynamically queries structure databases – at the time of writing this manuscript, variants were mapped onto 58,026 PDB structures of 7,973 proteins corresponding to 8,022 genes (39% of 20,292 genes) and 19,896 AlphaFold structures of 19,972 proteins corresponding to 20,026 genes (98.7% of 20,292 genes). Among 7,973 proteins with a PDB structure, 2,059 proteins (25.8%) have only one PDB structure reported, and the rest of the proteins (74.2%) have 14.5 PDB structures on average (**Supplementary Fig. 4** and **Supplementary Table 4**). AlphaFoldDB covers structures of 12,124 proteins whose experimental structure is not yet available. For those proteins, the median of the AlphaFold confidence score (pLDDT) for residue positions of variants is 90.4 (mean: 79.0), which demonstrates the utility of incorporating AlphaFold structures (**Supplementary Fig 5**). However, the predicted structure of a protein can be missing in the AlphaFoldDB when the protein is shorter than 16 or longer than 2700 amino acid residues, or the protein identifiers are newly added to UniProtKB and thus not yet included in the AlphaFoldDB. Out of 270 proteins missing AlphaFold structures, 106 have PDB structures, therefore, by retrieving both PDB and AF databases, G2P contains 3D structures of 20,131 genes (20,078 proteins) – 99.2% of 20,292 genes.

Approximately 95.4%, 90.0%, and 91.2% of gnomAD, ClinVar, and HGMD variants were mapped onto protein structures (see **Fig. 3a-c**), which are exportable in PyMOL-compatible format in the portal. The fraction of variants mapped onto PDB structures is higher for ClinVar and HGMD (35.5% and 49.4%, respectively) than for gnomAD (23.3%). The distribution of protein consequences and database-specific groups of variants (based on allele frequency for gnomAD, clinical significance for ClinVar, and confidence for HGMD) mapped on structures (**Fig. 3d-f**) illustrated that a higher fraction of ClinVar PLP variants could be mapped on PDB structures (14.5%) compared to AlphaFold structures (10.2%), reflecting the importance of experimental structures in pathogenicity interpretation. Similarly, for HGMD, the fraction of high-confidence disease mutations is higher in variants mapped on PDB (73.8%) than in variants on AlphaFold (67.0%). gnomAD variants do not show the difference in the distribution of allele frequency between variants on PDB and AlphaFold.

**Fig 3.**
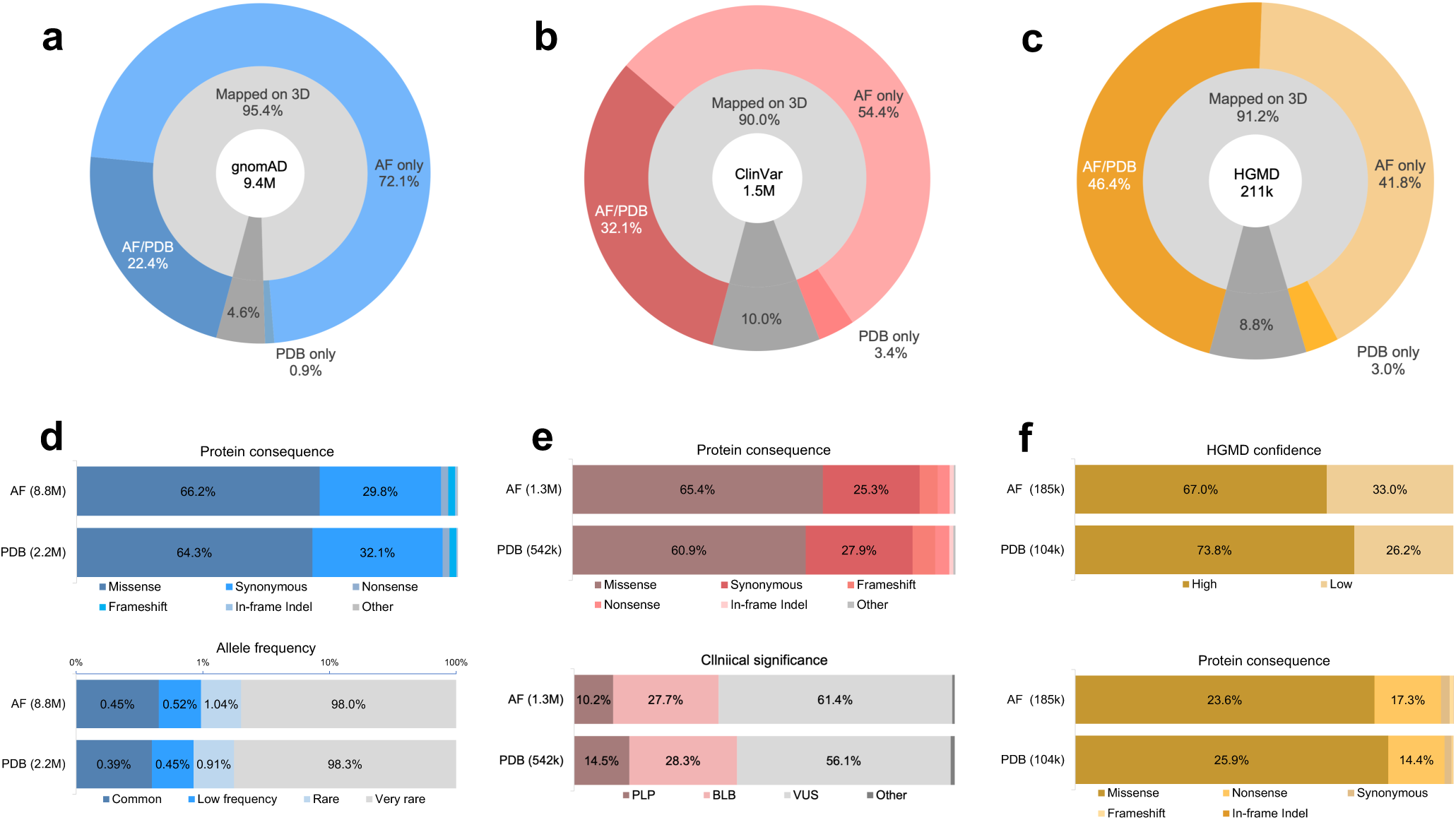
Statistics of variants mapped on 3D structures in the G2P portal. Variants annotated on transcripts corresponding to the canonical protein isoforms were mapped on 3D structures. The total number of canonical protein isoform variants from each database is shown in the middle of the donut chart. **(a)** The proportion of 9.4 million gnomAD variants mapped on PDB^8^, AlphaFoldDB^29^, or both. **(b)** The proportion of 1.5 million ClinVar variants mapped on PDB, AlphaFold, or both. **(c)** The proportion of 211k HGMD variants mapped on PDB, AlphaFold, or both. **(d)** The distribution of protein consequences (upper) and allele frequency group (lower) among gnomAD variants mapped on AlphaFold (8.8 million variants) and PDB (2.2 million variants). **(e)** The distribution of protein consequences (upper) and clinical significance (lower) among ClinVar variants mapped on AlphaFold (1.3 million variants) and PDB (542k variants). **(f**) The distribution of protein consequences (upper) and confidence (lower) among HGMD variants mapped on AlphaFold (185k variants) and PDB (104k variants).

#### Protein feature data

A comprehensive set of per-residue protein feature annotations have been integrated in the G2P portal, to help users establish the link between genetic variations and protein structure-function relationship. The features are grouped into (1) physicochemical properties of amino acids, (2) structural features^9,29,42,43^, (3) sequence annotation from UniProtKB^28^, (4) post-translational modification (PTM) information from PhosphoSitePlus^34^, and (5) readouts from multiplexed assays of variant effect when available in MaveDB^35^ (**Methods:** Protein features in the G2P portal).

G2P’s proteome-wide feature annotations provide insight into the characteristic structural and functional implications of disease-associated versus population variants. In **Fig 4**, we present the abundance and distribution of protein features across nine variant datasets, including missense variants in ClinVar (PLP, BLB, VUS), HGMD (*high* and *low* confidence), and gnomAD AF groups (*very rare, rare, low frequency,* and *common*). We observed a striking similarity in features between gnomAD variants with high allele frequency and ClinVar BLB variants, as well as between ClinVar PLP and HGMD high-confidence disease variants. **Fig. 4a** displays the abundance of UniProt sequence features and PTM sites (**Methods**: Normalized feature abundance calculation; **Supplementary Fig. 7**). Notably, ClinVar PLP and HGMD high-confidence missense variants exhibit a higher abundance of key functional sites (e.g., A*ctive site* and *Binding site*), biologically important domains or regions (*Domain, Repeat, Motif, etc.*), and PTM sites (*Acetylation, Methylation, etc.*) compared to gnomAD and ClinVar BLB variants, aligning with the expectation that variants perturbing protein function are likely to be pathogenic. Conversely, annotations related to disordered regions of the protein (such as *Region* and *Compositional bias*) were prevalent in gnomAD and ClinVar BLB missense variants.

**Fig. 4.**
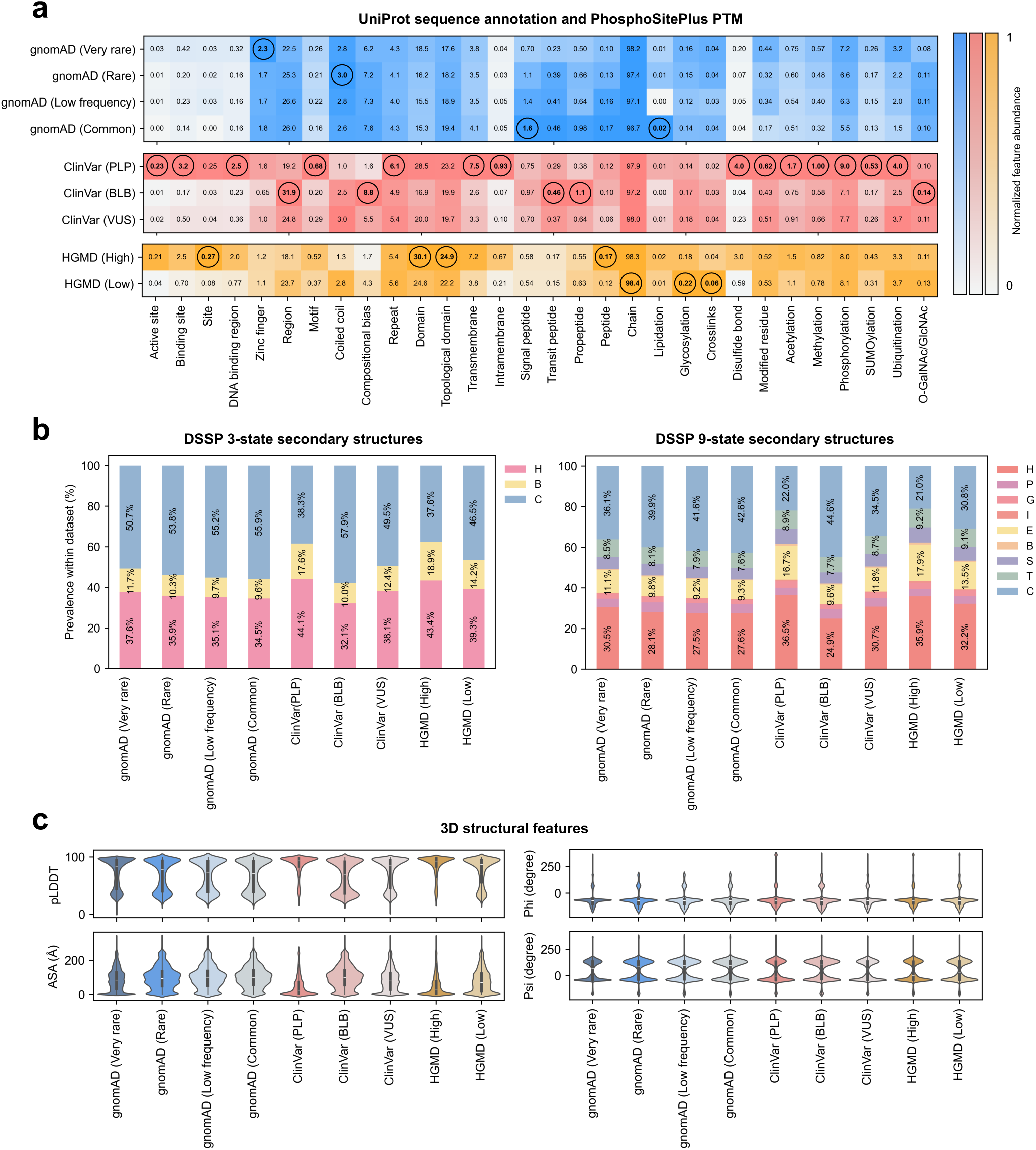
Abundance of protein features across nine variant datasets. These nine datasets include ClinVar variants grouped by clinical significance (PLP/BLB/VUS), HGMD disease mutations grouped by confidence levels (high/low), and gnomAD variants binned by allele frequency (*Very rare/Rare/Low frequency/Common*). **(a)** The abundance of each sequence annotation from UniProt and post-translational modification (PTM) site within a given dataset. The calculated abundance of a feature (e.g., *Active Site*) is denoted as the numerical value at each data point (see **Methods**: Normalized feature abundance calculation). Each point is color-coded based on its normalized abundance, wherein the abundance is divided by the maximum value among the nine datasets (denoted as bold and circled) to facilitate comparison of relative abundances across different features. For example, the abundance of the *Active Site* is the highest for the ClinVar PLP dataset, represented as 0.23, resulting in the darkest color where normalized abundance equals 1, while the gnomAD *Common* dataset has 0/23 = 0 having the brightest color. **(b)** The proportion of 3-class (*left*) and 9-class (*right*) secondary structures within variant datasets. Nine secondary structure classes are grouped into three larger classes: helix (H; 310-helix/G, α-helix/H, ρε-helix/I, and polyproline helix/P), strand (B; β-sheet/E and β-bridge/B), and loop (C; bend/S, turn/T, coil/C). Structured regions (helix and strand) have a higher prevalence of harboring pathogenic variants (62% of ClinVar PLP and HGMD High confidence disease mutations). **(c)** Violin plots for distributions of 3D structural features – AlphaFold pLDDT, accessible solvent area (ASA), backbone phi/psi angles. Notably, across all protein features, gnomAD Common and ClinVar BLB exhibit similar distributions of 3D features as well as ClinVar PLP and HGMD High.

The abundance of secondary structures and other 3D structural features indicates that unstructured protein regions tolerate mutations (**Fig. 4b-c**). gnomAD and ClinVar BLB variants showed a higher abundance of coils (C), bends (S), and turns (T), while ClinVar PLP and HGMD high-confidence variants exhibited a higher fraction of helix (H, P, G, I) and β-strand (B, E). This trend is also reflected in AlphaFold pLDDT, where pLDDT scores of gnomAD *low frequency* and c*ommon* variants distribute towards both high and low pLDDT values, while those of ClinVar PLP and HGMD high-confidence variants are concentrated around high-confidence regions. The distribution of accessible surface areas (ASA) of residues at variant positions highlights that high allele frequency gnomAD variants and ClinVar BLB variants have higher ASA, suggesting greater exposure to solvent (**Fig. 4c**). In contrast, ClinVar PLP and HGMD high-confidence variants show a localized peak at low ASA regions, indicating that missense variants at the protein core (low ASA) are likely to disrupt protein structures and lead to functional loss. Additionally, the frequency of substitution between 20 amino acids supports the similarity of occurrence of particular amino acid substitutions between benign/population datasets and between disease variant datasets (**Supplementary Fig. 8**). Investigation of protein feature abundance was repeated for other protein consequences (synonymous, nonsense, and frameshift) and results are given in **Supplementary Fig 9-10**.

### User interface of the G2P portal

The web-based user interface of the G2P portal (g2p.broadinstitute.org) is built in a streamlined and efficient Google Cloud infrastructure (**Methods:** G2P Google Cloud Infrastructure**, Supplementary Fig. 11)**. From the home page, users can access the Gene/Protein Lookup module by searching for a human gene or protein name. When available, users are directed to the protein sequence annotations (**Supplementary Fig. 12**). Upon selecting a transcript, users then can map genetic variants from gnomAD^14^, ClinVar^12^, and HGMD^13^ databases along with the protein features on the sequence. Subsequently, by selecting a structure from PDB and AlphaFold, users can transfer the variant and feature annotations to the structure. All human proteins and available structures (PDB^8^ and AlphaFoldDB^29^) annotated with genetic variants and protein features aggregated in the G2P portal are available for visual inspection and exportable via the Gene/Protein Lookup module of the portal.

Alternatively, from the home page, a user can access the Interactive Mapping module of the portal which allows users to securely upload genetic variant annotations at the protein residue level along with other protein sequence annotations, e.g., domains, drug-binding pockets, conservation scores, and map them to the target protein’s structure, essentially generalizing this capability of linking genomics to proteins beyond the genetic variants and protein structures available in the databases and in fact, beyond human proteome. For example, researchers can upload synthetic variations generated from functional genomics screens performed in a mouse model and map the results to the structure of the mouse protein predicted using a structure prediction method of their choice (RosettaFold^1^, ESMFold^3^, etc.). The data flow of the portal ensures the security of user-uploaded data (**Supplementary Fig. 11**); user-provided data are not shared or saved to the portal’s backend. Additionally, the versatile and interoperable pipeline of G2P allows concurrent mapping of multiple data types (genetic variants, discrete feature annotations, and continuous scores) from gene to protein sequence and structure and exporting the results for downstream analyses (see example use-cases below). Further details of submodules within the two main modules of the portal are available in **Methods**: G2P Portal Sitemap and **Supplementary Fig. 12**.

### Data visualization tools in the G2P portal

Across the two main modules of the portal (Gene/Protein Lookup and Interactive mapping; **Supplementary Fig. 12**), a suite of visualization tools has been implemented for intuitive exploration of the data – protein sequence viewer, variant information and protein feature cards, variant and protein feature tables, protein structure viewer, and mutagenesis output viewer.

#### Protein sequence viewer

We adopted the RCSB Saguaro 1D Feature Viewer^44^ and customized it for online visualization of variants and protein features mapped onto the protein sequence with dynamic applications of various filters, referred to henceforth as the protein sequence viewer (**Fig. 5a**). For example, mapping of *MORC2* ClinVar variants filtered by their pathogenicity shows that all pathogenic mutations are concentrated in the N-terminal (residues 20-470) of the protein. Tracks of features along with variants unveil notable observations – the N-terminal of *MORC2* has experimentally solved structures (PDBe/SIFTS track); the N-terminal have lower solvent accessibility and higher confidence in the predicted structure than the C-terminal (ASA and pLDDT track); it harbors important catalytic binding sites with ATP and ZN^2+^ (Binding sites track). In the protein sequence viewer, users can save their variant and feature mapping as a publication-ready figure. Additionally, users can view and download a protein residue-wise annotation of variants and details, and protein features in tabular format by clicking on “View as Table” (**Fig. 5a**; see **Methods:** Variant and protein feature table) and as PDFs. Clicking on a specific variant, users can access “Variant information card” and “Protein feature card” (**Supplemental Fig. 13;** see **Methods:** Variant and protein feature cards), summarizing details of the selected variants and protein features at the residue position.

**Fig. 5.**
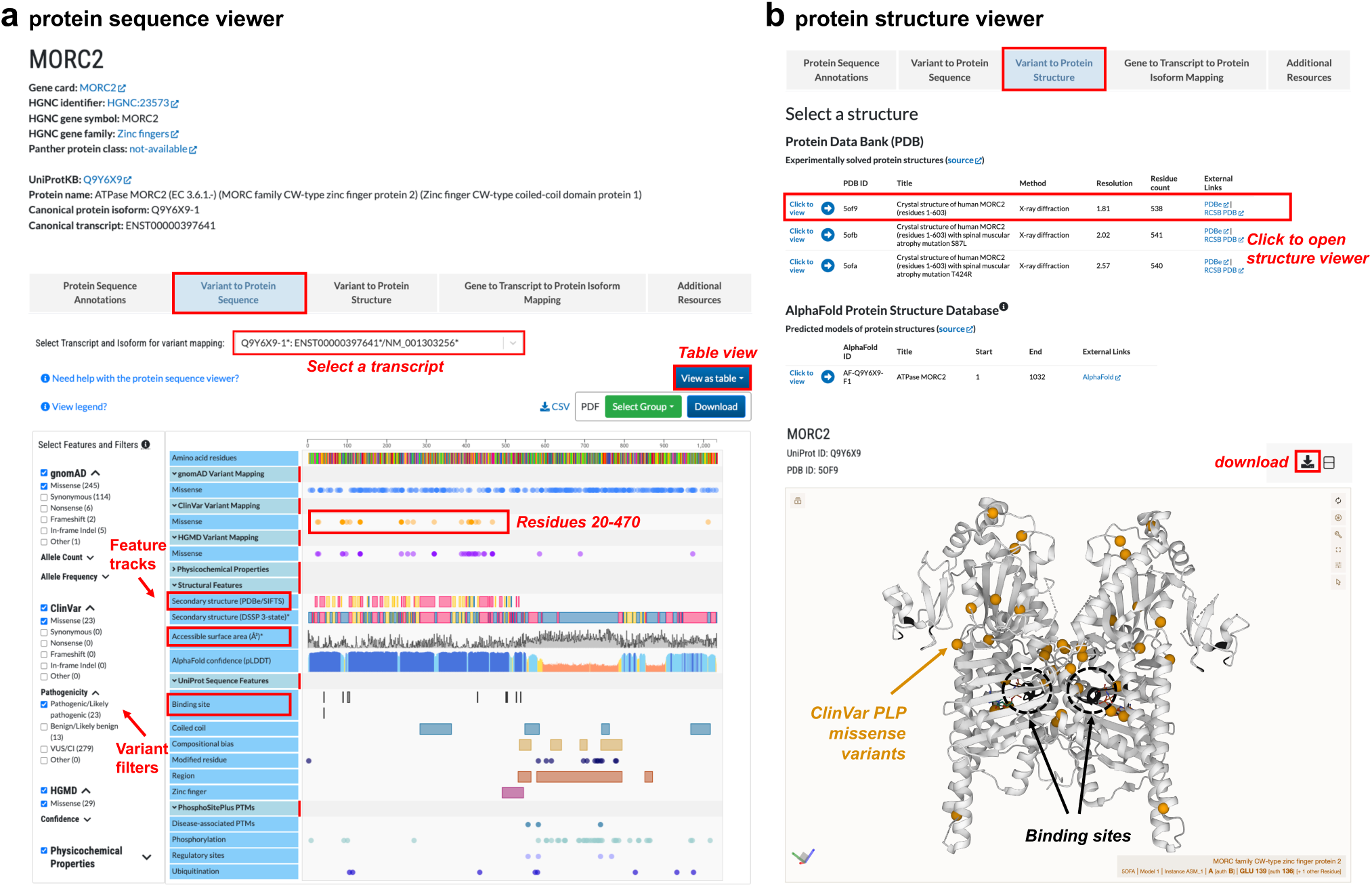
User interface and data visualization tools in the G2P portal. **(a)** Protein sequence viewer: the landing page of the Gene/protein lookup module shows an overview of the input gene information, followed by the protein sequence viewer displaying the aggregated protein features and variants for the target gene or protein. Protein features associated with the canonical protein isoform appear as default. Results can be seen and exported in tabular format (*View as table* button) and are also downloadable as PDFs (*Download* button). Variants can be filtered by categories (see **Results**: Resources in the G2P portal, **Fig. 2**). **(b)** Protein structure viewer: To map variants on a structure, users can navigate to *Variant to protein Structure* from the landing page of the Gene/protein lookup module, select a structure and “click to view”, which launches the protein structure viewer. For example, here, the viewer illustrates MORC2 structure (PDB: 5OFA) with concurrent mapping of ClinVar PLP variants track (yellow) and protein feature, *Binding site* (black). After performing the mapping of variants and features on the structure, users can download the results in a PyMOL-compatible file. The sequence and structure viewers are integrated in both Gene/Protein lookup and Interactive mapping modules of the portal.

#### Protein structure viewer

We integrated the Mol* protein structure viewer^45^ to visualize variants, protein features, and scores on protein structures, simultaneously with protein sequence (**Fig. 5b**). Users can map three types of data from sequence to structure: variants (mutation positions, as spheres), scores (continuous variable, as a heatmap), and multiclass features (discrete/categorical variable discretely colored by category). Users can map, review, and recolor features as desired, and apply data filters concurrently. For example, a user can filter ClinVar PLP variants and map them concurrently with UniProt sequence annotations as features (**Fig. 5b**), to identify pathogenic variants (orange spheres) around the binding sites (black). In the Interactive Mapping module, users can map user-uploaded annotations on the structure and can further add variant and feature annotations from available databases, to inspect user-uploaded data in the context of existing data.

The structure viewer is interconnected with the sequence viewer; when a user hovers over residues in sequence, they are highlighted in the structure, and vice-versa. The G2P portal is dynamically linked with and loads structures from the PDB and AlphaFoldDB. Many AlphaFold structures show high-confidence structured domains surrounded by low-confidence regions, which challenge users to analyze the structure by obscuring structured regions and globular domains. As such, the structure viewer provides additional functionality, allowing users to hide residues on AlphaFold structures based on the AlphaFold confidence of the structure (pLDDT). To export data for subsequent analysis, the structure viewer allows users to download structures and all accompanying features in a prepared PyMOL file, which includes user-uploaded and G2P-provided features as annotations in the PyMOL session.

#### Mutagenesis output viewer

To provide greater analytical capacity for integrated Multiplexed Assay of Variant Effect (MAVE) data, we implemented a 2D heatmap view of the data (when available, **Supplementary Table 5**), accessible via the *Additional resources* tab in the Gene/Protein lookup module (**Supplementary Fig. 12**). For single missense mutation MAVEs, variant scores are displayed as a 21x*N* heatmap, where *N* is the range of all perturbed residues, and the 21 rows correspond to the 20 possible amino acid mutations and the stop codon (**Supplemental Fig. 14**). For double mutation, scores are given as an *NxN* heatmap, where the row and column represent the first and second residue positions from a pair of jointly perturbed residues. Each entry corresponds to the average score of all perturbations in the MAVE at the corresponding pair of residue positions (see further details in **Methods**: MAVE output viewer).

### Case study – Interactive Mapping

**Fig. 6** presents a case study within the Interactive Mapping module by utilizing recently published base-editing (BE) scanning results of the DNA methyltransferase^19^. From “Start with a gene/protein identifier”, users can select a gene (*DNMT3A*), choose desired structures (*PDB: 4U7T*), and upload annotations (protein residues-wise variants, scores, and features). For this case study, we uploaded: (1) 34 missense variants (*Base-edited position*), with absolute sgDNA scores ≥ ± 2 standard deviation^19^, (2) *sgRNA* scores from the BE screen and the pathogenicity prediction scores from AlphaMissense^46^, and (3) domain annotations (*Domain*) provided in the paper^19^ (**Supplementary Table 6**). User-uploaded annotations are visible and selectable in the viewer (**Fig. 6a**) and users can supplement these with additional annotations from resources integrated in the portal (described in “Resources in the G2P portal”). By selecting *“Base-edited position”* and “*Domain”* annotations (**Fig. 6a**, *left*), the user can pinpoint the 3D position of variants within each domain (**Fig. 6a**, *right*) – 24 and 4 variants found in MTase and ADD domain, respectively. **Fig. 6b** illustrates the availability of concurrent mapping between user-uploaded base-edited variants and additional data, i.e., ClinVar PLP variants and 3-class secondary structures; this capability allows users to analyze their uploaded variants in the context of pathogenic variants (**Fig. 6b**, *top*) and structural features (**Fig. 6b**, *bottom*).

**Fig. 6.**
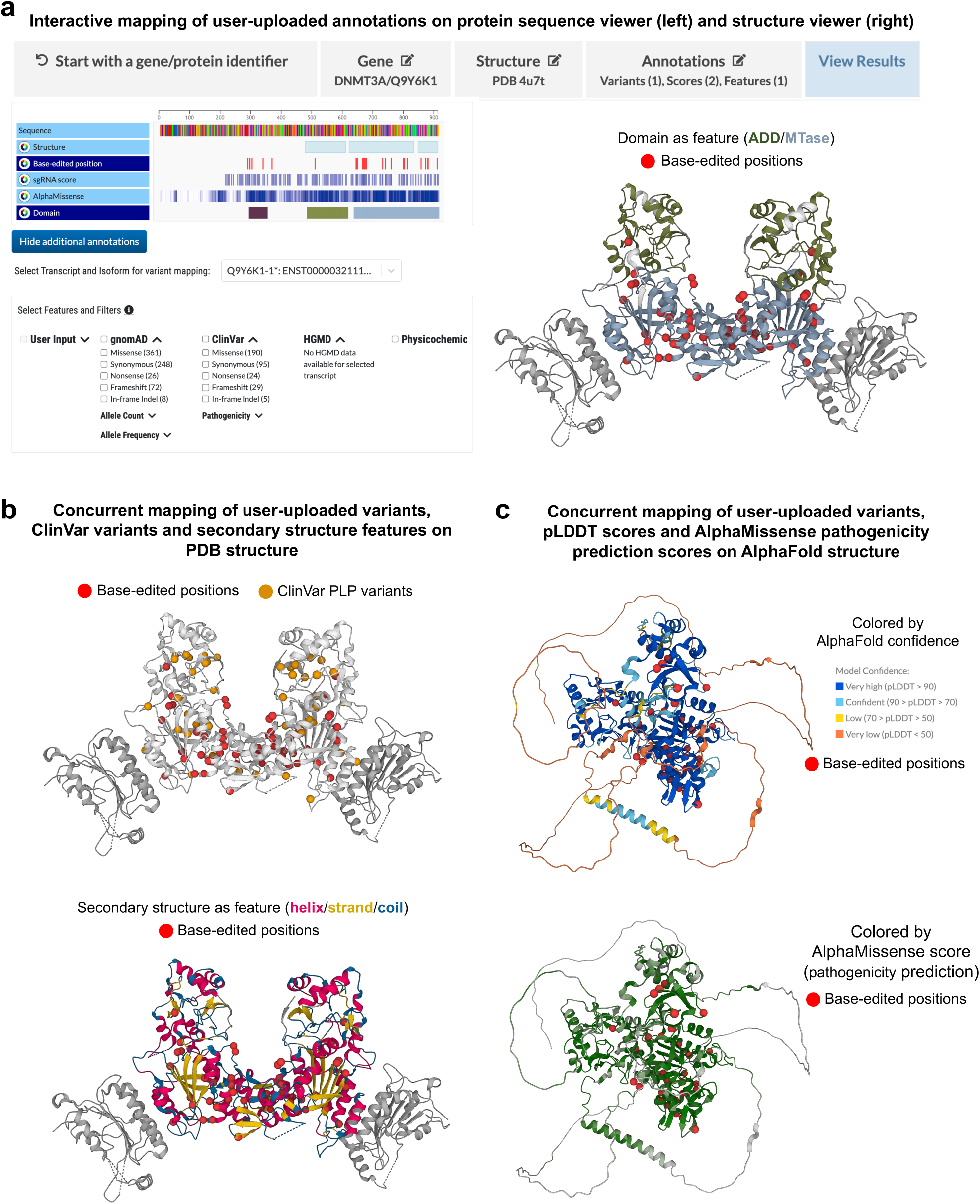
A use case of the Interactive Mapping module using *DNMT3A* base-editing screens. **(a)** The user interface of interactive mapping viewer. From “Start with a gene/protein identifier,” users are asked to select a gene (*DNMT3A*), structure (PDB: 4U7T), and upload annotations (variants, to be shown as spheres; continuous data or scores, to be shown as a heatmap; and discrete data or features, to be shown in discrete colors). The selected gene, structure, and entered annotations can be edited by going back through the workflow. Finally, in “View Results,” annotations are visible on sequence (*left*) and structure (*right*). The annotation tracks are selectable from the sequence viewer to map specific tracks on the structure. For example, the mapping on the structure viewer (right) is the result of clicking the “*Base-edited position”* and *“Domain”* tracks, where variation data are shown as *red* spheres and domain annotations are displayed as features in different colors. Colors are editable by the users. **(b)** Illustration of the concurrent mapping of user-uploaded variant annotations and data from additional G2P-provided resources on the structure (see **Results**: Resources in the G2P portal). On the upper panel, the *Base-edited positions* (*red* spheres) and the ClinVar PLP variants (*orange* spheres) are simultaneously mapped on the structure (MORC2, PDB: 7PFP). On the bottom panel, the *Base-edited positions* (*red* spheres) are displayed in the context of secondary structure annotations (as discrete features) available in the portal. **(c)** Illustration of the concurrent mapping of user-uploaded variants, features, and scores on the structure. The top panel shows the *Base-edited positions* (*red* spheres) in the context of pLDDT values (four discrete features: very high, confident, low, and very low)^29^, whereas the bottom panel shows the user-provided *Base-edited positions* (*red* spheres) in the context AlphaMissense^46^ pathogenicity scores (*green* spectrum) where darker green indicates higher pathogenicity scores. After performing a workflow in the Interactive mapping, users can download the current mappings as a TSV file (protein residue-wise annotation) and a PyMOL-compatible structure file.

Finally, **Fig. 6c** showcases the mapping of the BE result onto AlphaFold structures, colored by the predicted model confidence scores (pLDDT; upper panel) and AlphaMissense scores averaged over 20 amino acid substitutions at a given reference amino acid position (lower panel); a capability in the G2P portal that enable users to characterize variants using any state-of-the-art variant effect prediction score. In this case-study, we observed all 34 base-edited variants on residues with pLDDT > 70 and 31 variants (91% of 34) are pathogenic according to AlphaMissense (score > 0.57).

## Discussion

Genetic screening is increasingly applied in clinical practice, leading to the identification of a rapidly growing number of genetic variations^47–49^. One of the challenges in modern biology is decoding the functional implications of these genetic variations at the molecular level^37,38,40,50–55^. This is pivotal not only for unraveling the disease mechanisms^39,56–59^ but also for leveraging this knowledge to drive therapeutic developments^60–63^. Bridging genetic variants to structural biology provides a means to connect the potential cause of many genetic diseases to the molecular effect. However, integration of data across multi-omics is not a straightforward task due to different data types and inherent complexities. Genomics 2 Proteins portal is developed to address the challenge of connecting data across genomics, transcriptomics, protein sequence and structure, thereby, the tool has the potential to generate hypotheses on translating genetic discovery into molecular consequences and mechanisms.

A major bottleneck in translational and clinical genetics is that a large portion of clinically identified missense variants remains of uncertain significance^64,65^. Various in silico variant effect predictors (VEP) are currently available, providing a pathogenicity prediction score, generated using different classification algorithms, training cohorts, and features^42,46,66–70^. Early-stage VEPs such as SIFT^42^ or Polyphen^66^ relied on evolutionary conservation, followed by the utilization of large datasets encompassing population and clinical variants, as seen in tools like CADD^67^ or REVEL^68^. More recently, generative AI approaches have been adopted such as EVE^69^, ESM1v^70^, and AlphaMissense^46^. In addition to the prediction score, it would be beneficial for the biomedical community to be able to rationalize variant effects using biologically interpretable protein features^54,71,72^ and generate hypotheses in the context of protein structure and function to guide assays and drug design^41^. While recent efforts, including ProtVar^73^, offer functional annotations and some structural context of a variation location, the G2P portal integrates a comprehensive set of protein feature annotations per amino acid residues for the entire human proteome, providing insights into structural and functional impact of the mutation (**Fig. 4**). These encompass determining whether a variant is situated in the protein core/surface, on protein domain or disordered regions, or at the interface of a protein assembly, possibly leading to a dominant-negative effect^74^, etc. G2P’s concurrent mapping of population and disease variants along with the protein features can be utilized for identifying mutation-intolerant regions and prioritization of variants. Moreover, by integrating data from MaveDB^35^ (currently for 40 protein-coding human genes), G2P allows for analyzing functional readouts in the context of protein structures (**Fig. 5**) and we anticipate that this part of the resource will continue to expand as further datasets are generated and shared^75^. Besides visualizing variants and features on protein sequences and structures, users can export sequence annotations as text files for further data analyses and machine learning purposes as well as structure annotations as PYMOL-compatible files for deeper structural investigation.

It has been discussed before that AI-based structure prediction has limited application in pinpointing the impact of point mutations on structures^76^, however, it is conceivable that linking structural data with genomic and functional data will help understand the relationship between clinical data (i.e., pathogenic mutations) and their structural effect^77^. Towards this direction, the inclusion of the AlphaFoldDB in the G2P portal expanded the variant-to-structure mapping capability from 35% to 94%. Comparing the fraction of variants per gene mapped onto PDB structures versus high-confidence AlphaFold structures (pLDDT > 70), we observed that about 30% of genes benefited from increased ClinVar PLP and HGMD disease mutation mapping coverage enabled by AlphaFold (**Supplementary Fig. 6).** This emphasizes the significant role of accurate protein structure prediction in advancing our comprehension of the structural characteristics of disease-associated genes.

Furthermore, the growing landscape of natural and synthetic variants as well as predicted protein structures underscore the urgent, unmet need for a flexible, dynamic, and interactive tool for variant mapping on protein structures that go beyond existing databases. The Interactive Mapping module of the G2P portal allows for this – a capability distinct from existing tools. This module allows users to securely upload genetic variant annotations at the protein residue level along with other protein sequence annotations and map them over to the target protein’s structure to generate scientific hypotheses. For example, a clinician with a *de novo* mutation never reported before or a molecular biologist with a set of mutations out of a base-editor screen can upload their data by searching the target gene of interest and interactively link their data to the target protein’s sequence and structure, to further investigate the mutations at the protein level, as shown in the case study (**Fig. 6**), alongside already reported variants from population and clinical databases. Similarly, a structural biologist can upload a previously unsolved structure of a drug target or a structure model predicted by an AI-method^1–7^, and can map the known set of disease mutations onto the structure for structure-based rationalization of the impact of disease mutations.

In summary, the G2P portal is an open-source platform for human proteome-wide mapping of genetic variants to protein sequences and structures. The efficient and user-friendly web interface of the portal is built upon an integrated and dynamic bioinformatic method enabling rapid and easy investigation of genetic variants in the context of protein structure, which otherwise requires manual labor and is error-prone. Current limitations of the portal include the exclusion of non-canonical protein isoform variant mappings onto structures directly from Gene/Protein lookup module, which requires transferring of variant positions from non-canonical to canonical protein isoforms, as PDB structures are available for canonical isoforms only. However, users can generate structure models for non-canonical protein isoforms using AI methods and use the interactive module of the G2P portal to upload the predicted structure and perform variant mapping on the structure. We are committed to growing and maintaining G2P tools and resources. As a part of this expansion, we are incorporating cross-isoform and cross-species variant mappings, computational predictions of binding pockets^78^, or free energy change upon mutation^79,80^. The data and capabilities integrated into the portal will connect researchers across different fields of biology for a holistic understanding of how genetic variants impact protein structure and function and thereby facilitate the spectrum of basic biology research, from the translation of genetic discovery into better target selection to drug discovery.

## Data availability

All resources in this paper are available in the Genomics 2 Protein portal website (https://g2p.broadinstitute.org/). The G2P3D API is available at https://g2p.broadinstitute.org/api/gene/:geneName/protein/:UniProtAC/gene-transcript-map.

## Acknowledgements

This work was supported by the Merkin Institute for Transformative Technologies in Healthcare. We thank Alex Wagner and Guillaume Poncet-Montange for the scientific discussion, Behnoush Hajian for the scientific illustration of the G2P portal, and the PATTERN team at Broad Institute for the website design feedback. A.F.R. received funding from NIH/NHGRI grants UM1HG011969 and RM1HG010461. Contributions of A.F.R. in this work were supported by the Australian government.

## Author contributions

S.I. conceptualized the project and designed the study. S.K. performed the data analyses. J.S. D.N., D.H., and S.I. led the development of the G2P website. J.S., S.K., P.M., J.A.A., and A.F.R. contributed to the data curation. S.K., J.S., and S.I. wrote the manuscript. D.N., D.H., P.M., J.A.A., A.F.R., A.J.C, and A.B. reviewed the final manuscript. A.J.C., A.B. and S.I. contributed the funding acquisition, supervised the project and portal development.

## Online Methods

### Construction of G2P3D API

We have integrated public databases focusing on genes, transcripts, and proteins to build an API for seamless mapping of identifiers for genes, transcripts, protein sequences, and structures (API: G2P3D; **Fig 1**). HUGO Gene Nomenclature Committee (HGNC)^30^ maintains a curated online repository (https://www.genenames.org/) of approved genes along with their unique symbols and names for human loci. The Ensembl genome browser (http://useast.ensembl.org/)^31^ offers access to a wide range of genomic annotations. UniProt KnowledgeBase (UniProtKB; www.uniprot.org)^28^ provides the most current data on protein sequences and functions. These databases each specialize in different aspects of biology and are regularly updated; thus, there would be situations where gene symbols annotated in UniProtKB have been changed or withdrawn in HGNC, and UniProtKB IDs annotated in the Ensembl browser have been obsolete in the latest release of UniProtKB. To address this issue, G2P3D API has integrated UniProtKB, Ensembl, and HGNC, ensuring that it captures the most comprehensive and up-to-date information on genes, transcripts, and proteins.

First, we obtained a list of all human proteins from UniProtKB/Swiss-Prot (indexed by UniProt Accession or UniProtAC) and their corresponding HGNC IDs. Then, we retrieved gene symbols for each protein from HGNC with the provided HGNC ID. Subsequently, all Ensembl and RefSeq transcript identifiers along with corresponding UniProtKB protein isoform identifiers, were obtained via the Mart View API from Ensembl BioMart^81^ for the human reference genome GRCh38. These data were processed to map each gene symbol (HGNC) to its encoded UniProtKB/Swiss-Prot identifiers (UniProtAC) and then each protein isoform to its corresponding Ensembl and RefSeq transcript when available (Ensembl). Additional annotations were made for canonical protein isoforms, canonical Ensemble transcripts, and RefSeq MANE-select transcripts. Next, the Protein Data Bank (PDB)^8^ identifiers for the experimentally solved protein structures per protein were obtained using Graph-API (https://www.ebi.ac.uk/pdbe/graph-api/uniprot/unipdb/:UniProtAC) and the identifier for the predicted structure by AlphaFold^29^ was retrieved using API (https://alphafold.ebi.ac.uk/api/prediction/:UniProtAC). The G2P3D API, incorporated into the G2P portal (see example of the API output in **Fig. 1b**), links 20,292 HGNC genes (**Supplementary Table 1**) that encode 20,242 UniProtKB/Swiss-Prot human proteins corresponding to 42,413 protein isoforms, via 53,607 Ensembl transcripts and 57,543 RefSeq transcripts, to 77,923 3D protein structures (58,027 experimentally solved and 19,896 computationally predicted). The G2P3D API is available at https://g2p.broadinstitute.org/api/gene/:geneName/protein/:UniProtAC/gene-transcript-map.

### Variant and feature mapping onto proteins

Genetic variants are annotated at the transcript; for example, variants sourced from gnomAD^15^ are at Ensembl transcript (ENST-), and those from ClinVar^12^ are at RefSeq transcript (NM-). Each variant aggregated from the databases was linked to its corresponding protein isoform IDs using the in-house G2P3D API (**Fig. 1b**) and then mapped onto its amino acid position upon fetching the protein sequence using UniProt rest API (https://rest.uniprot.org/uniprotkb/UniProtAC.json). Variants were mapped to both canonical and non-canonical protein sequences but only to structures of canonical protein sequences. Users select the target gene, then the transcript and corresponding protein isoform, and the desired protein structure identifier for mapping variants and additional annotation from a gene to protein structures. Finally, proteins’ functional and structural features were annotated onto variant positions at the protein level (see **Methods: Protein Features in G2P portal** for details). The predicted structures covered the full-length protein sequences; however, the experimental structures often cover only parts of the protein and have gaps. We used the polymer_coverage API (https://www.ebi.ac.uk/pdbe/api/pdb/entry/polymer_coverage/) to map experimental structure coverage to the sequence space for each chain. We then map protein residue positions from sequence to structure and consequently transfer the variants (i.e., protein consequence positions) to protein structures, leveraging Mol*^45^ functionality to properly align variants to positions before and after gaps. We found some limitations with polymer_coverage API and Mol* coverage detection, for example, a gap in a PTEN structure, PDB: 5BUG, (i.e., missing region in crystallographic structure) is incorrectly reported in the API response and also incorrectly aligned in the Mol* software.

### Gene family and protein class annotations in the G2P portal

About 75% of genes (15,275 out of 20,292, **Supplementary Table 1**) were annotated to be a part of the HGNC^30^ gene family (834 gene families), while the rest have no family information available (**Supplementary Table 2**). On average, each gene family has 75 member genes. Some families with the largest number of members are zinc fingers (1562 genes), G protein-coupled receptors (829 genes), solute carriers (414 genes), etc. (**Supplementary Fig. 1a**). Additionally, the protein functional class annotations of 20,424 UniProtKB proteins corresponding to 20,292 genes were obtained from PANTHER knowledgebase (PANTHER release 18.0)^36^. About 69% of genes (13940 out of 20292) were annotated within 24 PANTHER protein classes (see **Supplementary Table 3**). Metabolite interconversion enzyme is the largest class, containing 1954 genes, while storage protein is the smallest class with 8 genes (**Supplementary Fig. 1b**).

### Variant aggregation

We downloaded raw VCF files (https://gnomad.broadinstitute.org/downloads/) for genome and exome datasets from gnomAD v2.1.1 and selectively extracted variants that passed all variant filters for quality control (filter=’PASS’ flag) and possessed valid HGVSp annotation. When the same variant was identified in both genome and exome datasets, we summed the allele count and the sample count, subsequently calculating the merged allele frequency value. Variant data from ClinVar (Nov 2023 release)^12^ were downloaded directly from the ftp site (https://ftp.ncbi.nlm.nih.gov/pub/clinvar/tab_delimited/variants_summary_txt.gz). Variants were filtered based on the reference genome GRCh38 and valid HGVSp annotation. From Human Gene Mutation Database (HGMD®) professional release (2023.1)^13^, variants on GRCh38 and with disease-causing state (variantType = ‘DM’ flag, indicating disease mutation), and with valid HGVSp annotation were extracted. Variants were excluded under the following conditions: (1) reference or altered amino acids are not 20 natural amino acids, and (2) the gene was not included from a list of genes from the G2P3D API, which contains only a reliable set of genes both present in HGNC and UniProtKB databases. The resulting variant aggregation spans 18,014,632 gnomAD variants, 1,749,628 ClinVar variants, and 232,183 HGMD variants mapped on protein sequences.

### Protein features in the G2P portal

The G2P portal provides a comprehensive set of protein features on both protein sequences and structures, which include physicochemical properties of amino acids, sequence annotations collected from external databases such as UniProtKB^28^ and PhosphoSitePlus^34^, 3D structural features collected from PDB^8^ and AlphaFold^29^, and readouts from the multiplexed assay of variant effect (MAVEs)^35^ when available in the MaveDB.

1. The physicochemical properties of reference amino acids. The 20 natural amino acids are grouped into 6 categories based on physicochemical properties of their side chain R-groups; (i) Aliphatic – alanine (Ala/A), isoleucine (Ile/I), leucine (Leu/L), methionine (Met/M), valine (Val/V); (ii) Aromatic – phenylalanine (Phe/F), tryptophan (Trp/W), tyrosine (Tyr/Y); (iii) Polar/Neutral– asparagine (Asn/N); glutamine (Gln/Q), serine (Ser/S), threonine (Thr/T); (iv) Positively-charged– arginine (Arg/R), histidine (His/H), lysine (Lys/K); (v) Negatively-charged– aspartic acid (Asp/D), glutamic acid (Glu/E); (vi) Special – proline (Pro/P; a cyclic side chain and cannot make backbone hydrogen bonds), glycine (Gly/G; does not have a side chain that allows flexibility) cysteine (Cys/C; a reactive sulfhydryl group -SH in the side chain). In addition to these groupings, the molar mass (g/mol) and hydropathy index (a numerical measure reflecting the hydrophobicity of a side chain – the larger the number is, the more hydrophobic the amino acid) of each amino acid are shown for the protein sequence.
2. 3D structural features; The G2P portal provides pre-computed annotations on structural features. These features are computed based on AlphaFold predicted structures, aiming for extensive coverage. Secondary structures of amino acids refer to the local three-dimensional (3D) conformations of the polypeptide backbone. DSSP (Define Secondary Structure of Protein) is the standard tool for determining secondary structure by classifying each residue into a 3-class structure (H; helix, B: β-sheet/strand, and C; loop/coil) or a 9-class structure (G; 310-heilx, H; α-helix, I: π-helix, P; polyproline helix, B: isolated β-bridge, E; parallel β-sheet, S; bend, T; turn, C: loop/coils). We utilized DSSP to annotate both 3-class and 9-class secondary structures on AlphaFold2 structures. When experimental structures are available (e.g. from PDBe/SIFTS), we provide PDBe/SIFT secondary structures, which are derived from experimental structures, in a separate track. Additionally, DSSP calculates the accessible surface area (ASA in Å2) and the backbone torsional phi/psi angles (in degrees) for each amino acid position within the context of the protein’s 3D structures. Furthermore, we include a per-residue confidence score produced in AlphaFold, known as pLDDT (predicted Local Distance Difference Test). The score ranges from 0 to 100 and categorizes the confidence as “Very high” (pLDDT > 90), “High” (pLDDT > 70), “Low” (pLDDT > 50), or “Very low” (pLDDT < 50). Residues are color-coded accordingly. It’s important to note that residues with very low pLDDT may indicate that their structures are disordered in isolation.
3. Sequence annotation from UniProtKB: UniProt database was mined to gather the sequence annotations that describe various regions, domains, or sites of interest for a protein, elucidating its function, binding, sequence motif, domain/site/region, molecular preprocessing, and more. The G2P portal offers 31 selected sequence annotations: Active Site, Binding site, Chain, Coiled coil, Compositional bias, Cross-link, Disulfide bond, DNA binding, Domain, Glycosylation, Initiator methionine, Intramembrane, Lipidation, Modified residue, Motif, Mutagenesis, Non-adjacent residues, Non-standard residue, Non-terminal residue, Peptide, Propeptide, Region, Repeat, Sequence conflict, Sequence uncertainty, Signal, Site, Topological domain, Transit peptide, Transmembrane, Zinc finger.
4. Post-translational modification (PTM): PTM refers to the covalent and enzyme-mediated modification of proteins to form mature proteins. We collected amino acid positions of six different PTM types from the PhosphoSitePlus database: (i) acetylation – introduces an acetyl group; (ii) methylation – addition of a methyl group; (iii) O.GlcNAc – also known as O-linked N-acetylglucosamine, is a form of protein glycosylation; (iv) phosphorylation – attachment of a phosphoryl group; (v) SUMOylation – addition of SUMO (small ubiquitin-like modifiers) molecule; (vi) ubiquitination – attachment of ubiquitin.
5. Readouts from multiplexed assays of variant effect (MAVE): MaveDB is a public repository dedicated to housing datasets from Multiplexed Assays of Variant Effects (MAVEs). These datasets primarily result from deep mutational scanning or massively parallel reporter assay experiments. When a gene/protein is available in MaveDB, amino acid positions displaying variants whose effect falls within the top and bottom 99th percentile are visibly highlighted in the sequence annotation viewer. For an in-depth exploration of a specific MaveDB entry, the entire datasets are displayed in the *Additional Resource* tab (**Supplementary Fig. 12**) in a visually informative format through a heatmap of the variant effect index. MAVE scores are downloadable in JSON format.

### Normalized feature abundance calculation

Feature abundance is defined as the percentage of variants annotated with a specific feature among the total variants. The calculation is performed for nine variant datasets: ClinVar (PLP, BLB, VUS), HGMD (high and low confidence), and gnomAD AF groups (very rare, rare, low frequency, and common).

The abundance values calculated from these datasets vary due to differences in the natural occurrence of features. **Supplemental Fig. 7** illustrates the computation of normalized feature abundance using the example case “Active site.” For instance, *Active site* annotation ranges from 0 to 0.23%, while the *Chain* annotation ranges from 96.7% to 98.4%. Therefore, we normalized the calculated abundance by the maximum value among the nine datasets (indicated in bold font in **Fig. 4a**). This normalization ensures that a dataset with the largest or smallest abundance of a feature appears as the darkest or brightest color-coded, respectively.

### G2P Google Cloud Infrastructure

The schematic view of the G2P portal infrastructure is presented in **Supplementary Fig. 11**. The G2P platform consists of a React.js web app, which is served by a Node.js backend running on Google Cloud Platform (GCP). The RCSB Saguaro 1D Feature Viewer^44^ and Mol*^45^ are adopted and customized as protein sequence and structure viewer, respectively, to visualize the frontend data on protein sequences and structures. The backend runs on Google App Engine, which is a serverless and on-demand compute offering that launches a variable number of backend instances proportional to usage.

Google Cloud Storage (GCS) is utilized as the primary data store for variant and feature annotations per gene alongside an in-memory datastore used on the G2P backend to track gene mapping relationships. The static data stored in GCS is collected, processed, formatted, and uploaded by the portal admin (**Supplementary Fig. 11**). To load static data from GCS, the portal requests files directly from the frontend, which reduces latency by avoiding an additional “hop” where data must first travel to the backend before reaching the frontend. From our testing, the minimum observed time for a backend request is 60ms round trip, and by requesting files directly from the frontend, G2P portal saves a minimum of 60ms per request. To load data from the in-memory datastore, the portal frontend makes requests to backend APIs, and the backend retrieves and returns the relevant records. The datastore is managed directly by the backend server, not by a separate process. In addition to managed data sources, the portal dynamically requests data from external APIs to provide the most current information possible (**Supplementary Fig. 11**). To this end, the G2P portal web app requests the latest protein sequence and structure records directly from UniProt, PDBe, AlphaFold, and EMBL-EBI APIs. In the Interactive Mapping module of the portal, users can provide their own data (protein residue-wise annotation of variants, features, scores, and protein structures) for joint analysis of user data with G2P-provided resources (see **Results**: Resources in the G2P portal). The interactive mapping module can be securely accessed via Google sign in, and to further secure data confidentiality, all user-uploaded data remain on the user’s browser only; therefore, no user-provided data leaves the user’s device. This ensures that the user has full, secure control over their data while simultaneously providing access to G2P Portal’s variants and protein features for joint analysis.

When a user selects a Gene/Protein via the Gene/Protein Lookup or as part of the Interactive Mapping workflow, static mapping information is fetched directly with the G2P3D API to connect Gene to Protein to transcript to sequence to structure. Subsequently, detailed gene and protein-specific data is fetched as static data from Google Cloud Storage and dynamic data from external APIs.

### G2P portal sitemap

The portal’s homepage is the central hub for navigating two primary modules, complemented by a top navigation bar featuring tabs for About, Documentation, and Feedback (**Supplementary Fig. 12**).

The Gene/Protein lookup module empowers users to search for a human gene or protein of interest, leading them to a comprehensive overview page containing five submodule tabs for further exploration. *Protein Sequence Annotations* tab hosts a protein sequence viewer, allowing users to choose a protein isoform identifier from the available UniProtKB list. The protein sequence viewer displays a complete list of protein features aggregated within the G2P portal (**Method**: Protein features in the G2P portal). Data can be presented in table view and is downloadable in TSV or PDF format. The *Variant to Protein Sequence* tab permits users to select an RNA transcript ID, adding variant data from gnomAD^15^, ClinVar^12^, and HGMD^13^ on the top of the sequence viewer (**Fig. 5a** and **Supplementary Fig. 12**). The sequence viewer, including variants, can also be exported to the table view, available in TSV and PDF formats. Clicking on a specific variant within the feature viewer generates a summary card with detailed information on the variant and protein features at the variant position (**Methods:** Variant and protein feature cards; **Fig. 5c**). Clicking on the *Variant to Protein Structure* tab, users can choose a structure from the available PDB and AlphaFold structures (**Fig. 5b** and **Supplementary Fig. 12**). The structure viewer, coupled with the sequence viewer, facilitates dynamic feature selection and variant filters. Outputs from the structure viewer are exportable in PyMOL-compatible formats. *Gene to Transcript to Protein Isoform Mapping* tab provides a table view illustrating the mapping between gene, transcript, and protein sequences, downloadable in TSV format. *Additional Resources* tab offers external gene information links, such as USCS^82^, ChEMBL^83^, DrugBank^84^, Orphanet^85^, and OMIM^86^. Furthermore, the portal integrates multiplexed assays of variant effect (MAVE) data from MaveDB for 40 genes^35^ (**Supplementary Table 5**), presenting mutagenesis scores through 2D heatmaps (**Fig. 5d**). Raw JSON files for scores are available for download.

In the Interactive Mapping modules, users can start their exploration from either a gene/protein identifier or their own protein structures. When starting with a gene/protein identifier, the portal retrieves sequences from the UniProt sequence API and available structures for user selection. Alternatively, users starting with their own protein structures can upload them in PDB format. In both scenarios, the final step prompts a window for annotations, providing a sample format and allowing users to enter their annotations (variants, scores, or features). The resulting data is displayed in the *View Results* section (**Fig. 6a** and **Supplementary Fig. 12**), featuring both sequence and structure viewers. When starting with a gene/protein identifier, users also can append additional feature annotations, such as protein features and variants, corresponding to the selected transcript or protein isoforms, and map them simultaneously with the user-uploaded data on protein sequences and structures (**Fig. 6a**).

### Variant and protein feature table

Users can view per-residue annotation of variants and protein features aggregated in the G2P resources from the portal. For gnomAD variants, the table include the HGVSp, HGVSc, Allele Count, Allele Frequency, Allele Number, Male Count, and Hemizygote count (Example: https://g2p.broadinstitute.org/table/LDLR/P01130-1/ENST00000558518/missense) For ClinVar variants, the details include Genomic Consequence, Protein Consequence, Variant Type, Clinical Significance, Phenotypes, the last evaluated data, and review status (Example: https://g2p.broadinstitute.org/clinvartable/LDLR/P01130-1/ENST00000558518/clinvar_single). For HGMD variants, the table include the transcript, HGVSc, Codon change, HGMD confidence, and HGMD disease (Example: https://g2p.broadinstitute.org/hgmdtable/LDLR/P01130-1/ENST00000558518/missense). The protein feature table (e.g., https://g2p.broadinstitute.org/features/LDLR/P01130/P01130-1) includes all features described in **Methods**: Protein features in the G2P portal. Data in these tabular views can be downloaded as machine-readable text files for further usage by users.

### Variant and protein feature cards

After mapping variants on protein sequences via the Gene/protein lookup module of the portal (**Fig. 5a**), users can click on a variant position to view detailed variant information for the selected variant (Variant information card, **Fig. 5b**), alongside residue-specific protein feature annotations (Protein feature card). For gnomAD, variant details include allele count, allele frequency, allele number, male count, and hemizygote count across all samples, as well as allele count and allele frequency for exome, genome, and populations. For example, in the case presented in **Fig. 5a**, users can select the *MORC2* ClinVar missense variant at Ser87, and a card will display below revealing details for variants Ser87Leu and Ser87Pro, along with structural and functional annotations on Ser87 (**Fig 5b**). For ClinVar, variant details include clinical significance, phenotypes, review status, and the last evaluated date. For HGMD, variant details include HGMD confidence and HGMD disease. Alongside the variant details, residue annotation data includes physicochemical properties, including chemical polarity, molar mass, and hydropathy; structural features, including SIFTS^42^ secondary structure from experimental structures, DSSP^43^ 3-state and 9-state secondary structures, phi/psi angles, accessible surface area, and pLDDT for AlphaFold predicted structures^29^]; UniProt sequence features^28^ including domain, disordered region, binding site, repeat, zinc finger, compositional bias, motif, and transmembrane annotations; PhosphoSitePlus PTM annotations^34^ including phosphorylation, acetylation, methylation, ubiquitination, and glycosylation; and MAVE scores from the top and bottom 95^th^ percentile of scores from MaveDB^35^. For MAVEs, the feature card includes a download link for a JSON download of all MAVE scores for a given residue position.

### MAVE output viewer

In the Additional Resources tab of the Gene/Protein Lookup module (**Supplementary Fig. 12**), users can view MaveDB^35^ information from different MAVE datasets in expanded detail. For single missense mutations, a *21xN* heatmap is displayed, where N is the range of mutations covered by MAVE perturbations with 21 rows for the 20 different amino acids and the stop codon possible at each position. Each value in the heatmap corresponds to the score recorded in the MAVE, or the average of multiple scores if multiple scores were recorded for the same mutation. An example is shown in **Supplemental Fig. 14** for *CBS* MAVE readouts collected via DMS-TileSeq at low levels of Vitamin B6. Scores show a clear distinction between residues 90-390 (low scores in blue) and residues at the N and C-terminus (high scores in red). For double mutant MAVEs, where 2 different residues were perturbed concurrently, an *NxN* heatmap is displayed where the row and column each represent one of the 2 residue positions perturbed in the experiment. As with the single missense mutations, the value in the heatmap corresponds to the reported score from the mutation or the average of all scores reported for the residue pair. Different MAVEs utilizing different techniques have different score scales and scores that require interpretation in the context of the methodology used by the corresponding MAVE. To this end, the G2P portal includes a brief description of the experimental technique and scoring methodology of the paper, as provided by MaveDB, and additional links to the score set page in MaveDB and the associated publication such that users can best understand the experimental conditions under which any specific score of interest was collected. To facilitate deeper analysis, the portal includes a downloadable JSON file with all coding and non-coding variants from the MAVE.

