## Supplemental Fig. for "Genomics 2 Proteins portal: A resource and discovery tool for linking genetic screening outputs to protein sequences and structures"

### Supplemental Figures

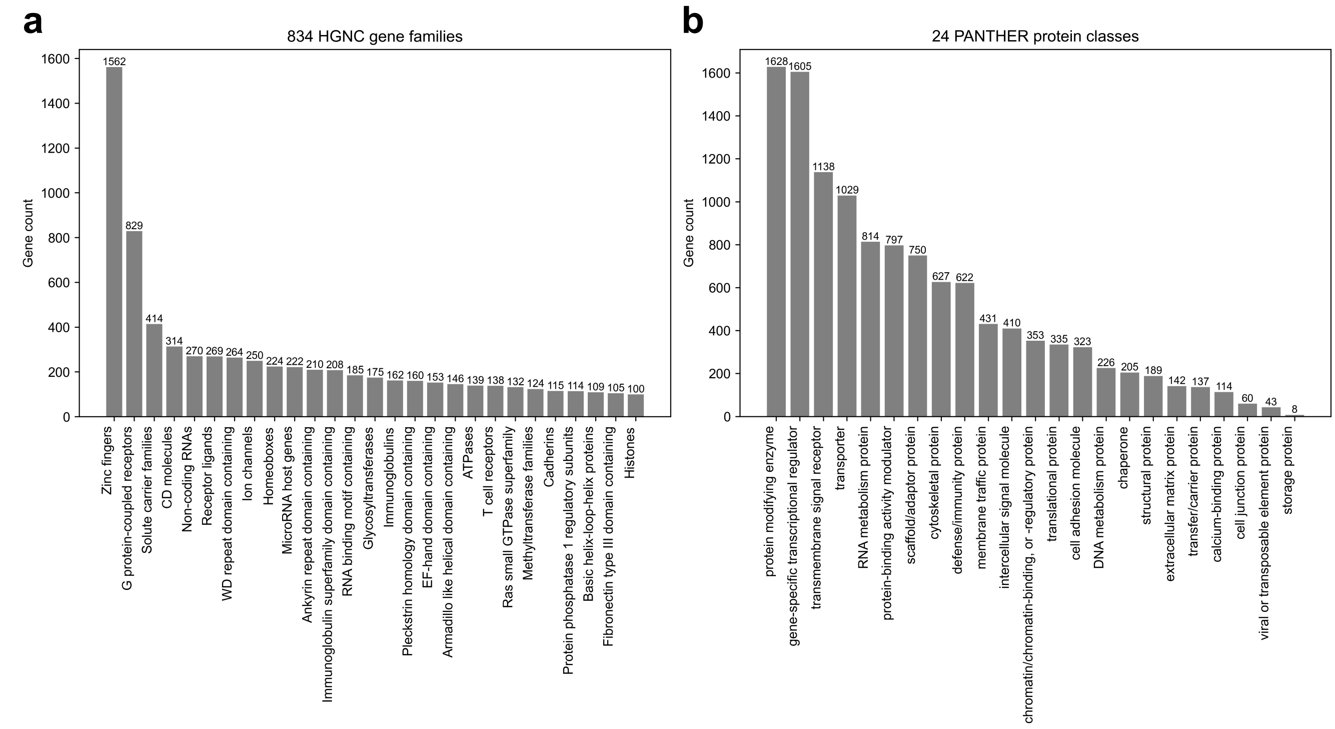


**Supplemental Fig. 1| Count of genes within HGNC family and PANTHER classes. (a)** The gene count within an HGNC gene family. Data are presented for families having over 100 genes, and the full data is available in **Supplemental Table 2**. **(b)** The gene count within 24 PANTHER protein classes. The full data is available in **Supplemental Table 3.**

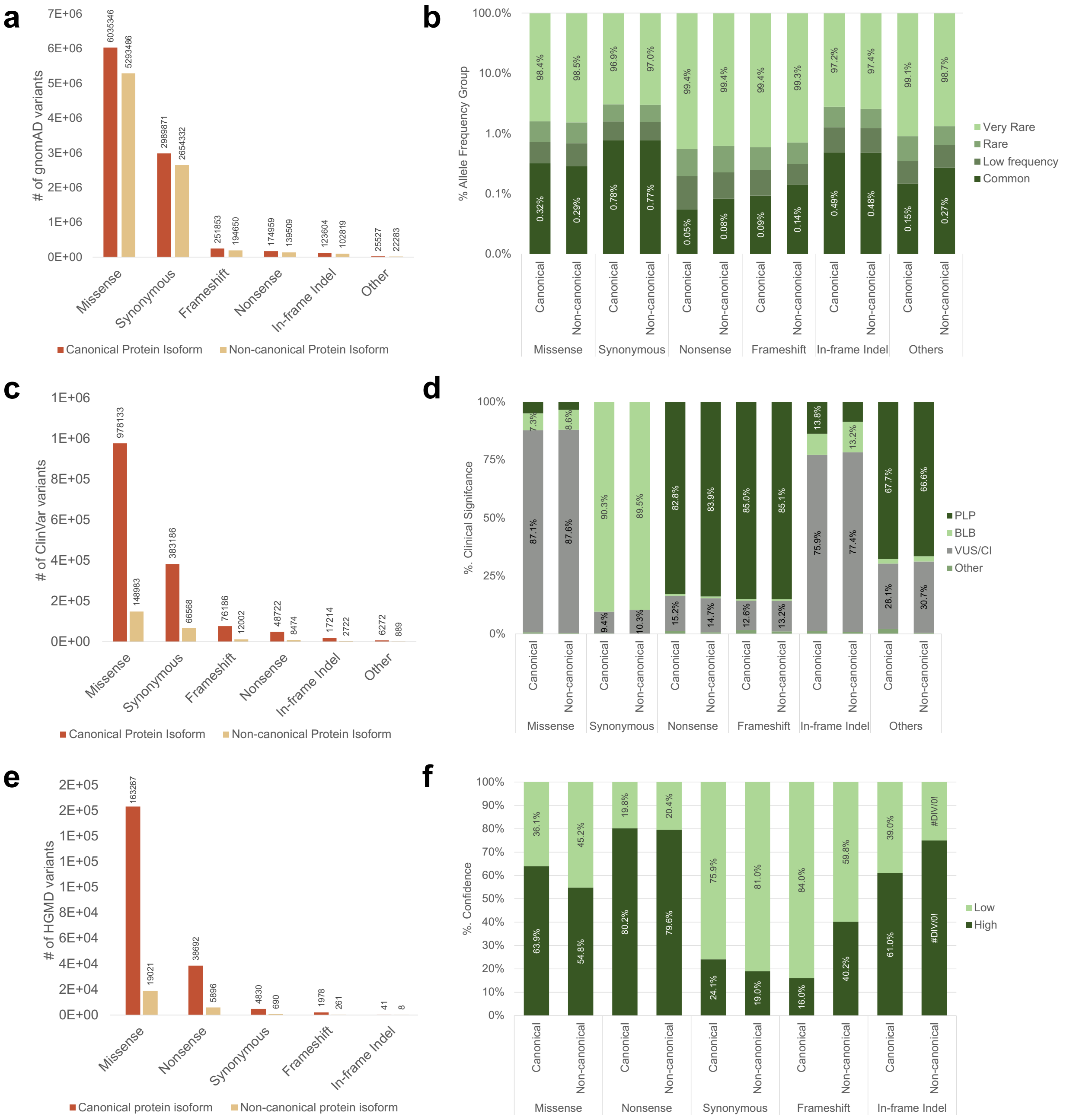


**Supplemental Fig. 2| Variant statistics on canonical vs non-canonical protein isoforms.** Variants were mapped on transcripts either on transcript coding canonical or non-canonical protein isoforms. The number of variants mapped on canonical and non-canonical protein isoforms are plotted across different protein consequences in **(a)** gnomAD, **(b)** ClinVar, and **(c)** HGMD databases. In addition, distributions of database-specific groups: **(d)** allele frequency for gnomAD, **(e)** clinical significance for ClinVar, and **(f)** confidence for HGMD are shown. Overall, the difference in statistics between canonical and non-canonical isoforms is marginal.

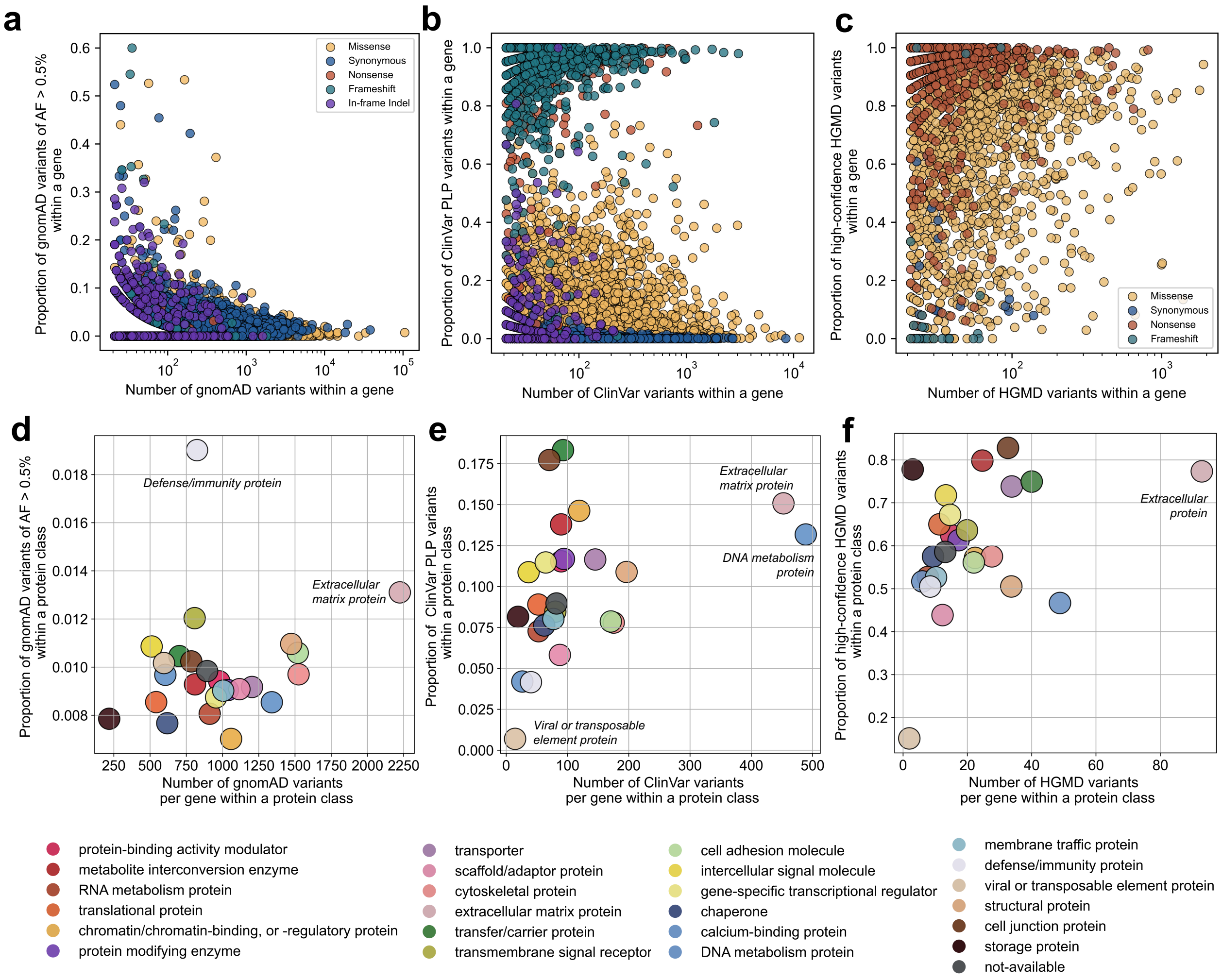


**Supplemental Fig. 3|** Gene-wise relationship between **(a)** the number of gnomAD variants and the fraction of gnomAD common variants of different protein consequences, **(b)** the number of ClinVar variants and the fraction of ClinVar PLP variants of different protein consequences, and **(c)** the number of HGMD variants and the fraction of HGMD high-confidence variants of different protein consequences. Each point in the scatter plots (a)-(c) represents a gene. Genes in the upper left quadrant of scatter plots (a) signify a lower overall variant count, with a notable proportion exhibiting high allele frequency (exceeding 0.5%), primarily due to synonymous variants. Genes associated with frameshift and nonsense variants tend to cluster at high proportions of ClinVar PLP or high-confidence HGMD variants, whereas genes linked to synonymous variants are predominantly at low proportions, and missense variants are widely distributed across varying pathogenicity. Similar analyses are done for different PANTHER protein classes: the scattered plot of 24 protein classes represented as different colors for **(d)** gnomAD, **(e)** ClinVar, and **(f)** HGMD, respectively. The x-axis is the number of variants per gene within a specific protein class. Some protein classes (e.g. *Extracellular protein* and *DNA metabolism protein)* have a higher number of population (gnomAD) or disease-associated (ClinVar and HGMD) variants, while some have lower (e.g. *Viral or transposable protein*). *Defense/immunity protein* have a moderate number of gnomAD variants (~800 per gene), but the proportion of high allele frequency (>0.5%) is the largest among all protein classes.

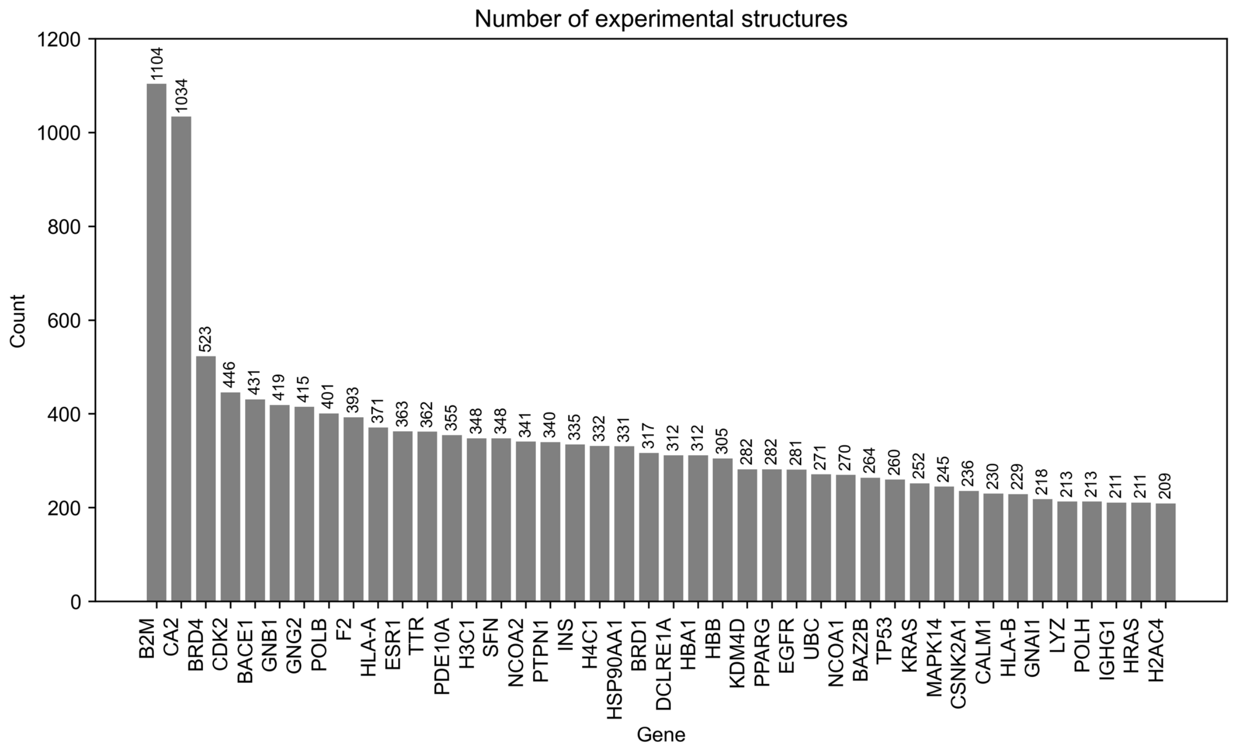


**Supplemental Fig. 4 | PDB Structure Counts**. This figure depicts the quantity of PDB structures available for each gene, sorted by the respective counts. The statistics are current as of the paper writing date (11/2023) and are expected to increase over time.

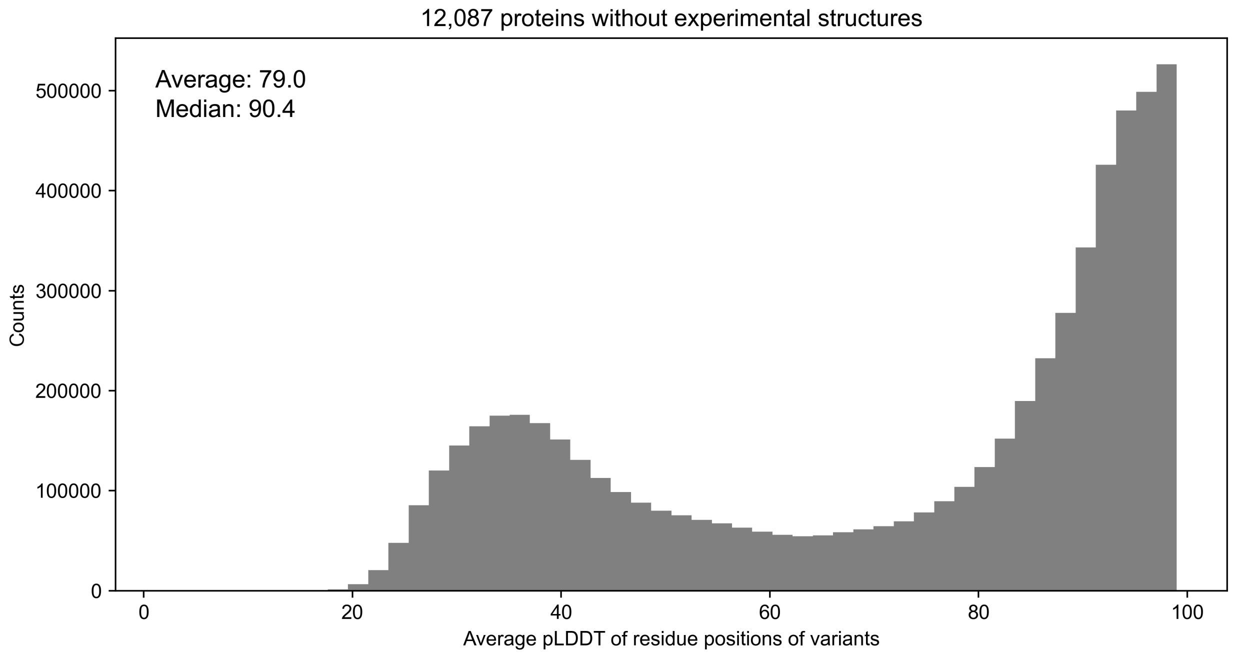


**Supplemental Fig. 5| The distribution of pLDDT of residue positions.** The figure illustrates the distribution of AlphaFold pLDDT scores of residue positions of variants (all gnomAD, ClinVar, and HGMD) found in 12,078 genes lacking available PDB structures (these variants constitute the proportion labled as “AF only” in Fig. 4(a)-(c)). The pLDDT distribution reveals two peaks at low and high pLDDT representing disordered and high-confident structure regions of predicted structures, respectively. The average and median pLDDT values are 71.2 and 82.9, respectively, affirming the reliability of utilizing AlphaFold structures for protein 3D feature analysis.

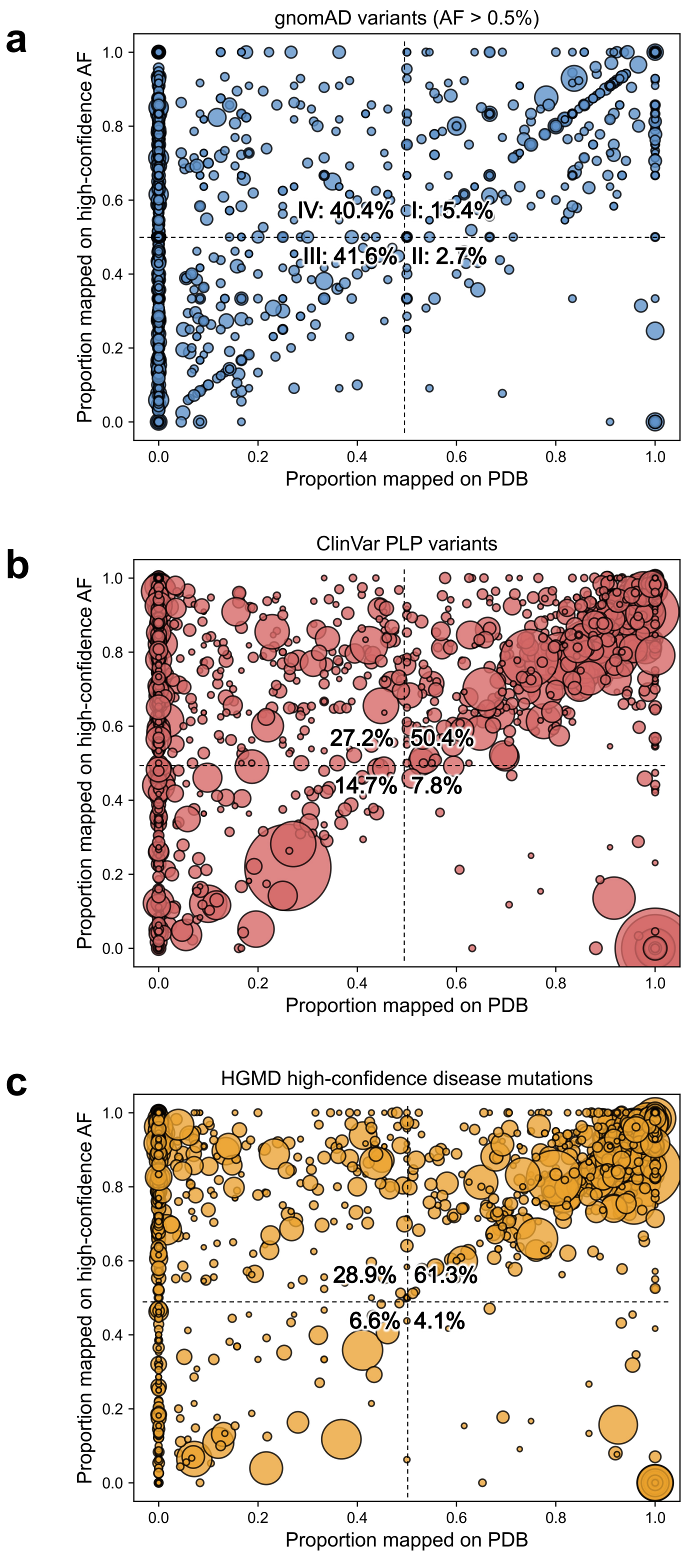


**Supplemental Fig. 6| Scatter plot of gene distribution of the proportion of variants mapped on the Protein Data Bank (PDB) compared to high-confidence (pLDDT > 70) AlphaFold structures.** We found 1.3M ClinVar,

192k HGMD, and 6.8M gnomAD variants were successfully mapped on protein structures. We calculated the proportion of variants mapped on the PDB and high-confidence AlphaFold structures within a given gene. For example, *LDLR* has 821 gnomAD variants on canonical protein isoforms, 689 mapped on PDB structures and 821 mapped on AlphaFold structures. Therefore, the proportion of variants mapped on PDB and AlphaFold is 84% and 100%, respectively. This analysis is applied to **(a)** gnomAD variants of AF > 0.5%, **(b)** ClinVar PLP variants, and **(c)** high-confidence mutations from HGMD. Each marker represents a gene, and the size of the marker indicates the total number of variants mapped on 3D structures. The graph is partitioned into four regions (I-IV), offering insights into genes with varying PDB and AF coverage. For instance, genes in Region I demonstrate comprehensive coverage by both PDB and AlphaFold, while those in Region IV exhibit limited coverage by both resources. Notably, the fraction in Region IV signifies the proportion of genes effectively rescued by AlphaFold, enabling the mapping of variants in genes lacking experimentally determined structures onto computationally predicted ones.

**Calculation of normalized feature abundance in variants** (example: Active sites in missense variants)

| Dataset | Number of variants | Number of variants annotated as ”Active Site” | Abundance (%) | Normalized abundance |
|---|---|---|---|---|
| gnomAD (Very rare) | 591460 | 1496 | 0.025 | 0.11 |
| gnomAD (Rare) | 52385 | 3 | 0.0057 | 0.02 |
| gnomAD (Low frequency) | 24508 | 2 | 0.0082 | 0.03 |
| gnomAD (Common) | 18784 | 0 | 0 | 0 |
| ClinVar (PLP) | 48487 | 111 | **0.23** | **1** |
| ClinVar (BLB) | 78897 | 4 | 0.0055 | 0.024 |
| ClinVar (VUS) | 85876 | 182 | 0.021 | 0.093 |
| HGMD (High) | 105136 | 225 | 0.21 | 0.93 |
| HGMD (Low) | 59525 | 21 | 0.035 | 0.15 |

**Supplemental Fig. 7| Calculation of normalized feature abundance.** This is an example of the calculation of normalized abundance of “Active site” UniProtKB sequence annotation in nine missense variant datasets (1st column). The abundance is the percentage of variants annotated “Active site” (3rd column) out of the total number variant of variants (2nd column). Abundance values (4th column) from nine different datasets are normalized by the maximum value, i.e., 0.23 in ClinVar (PLP) for “Active Site”, to compute normalized abundance (5th column). The normalized feature abundance is computed and shown for each sequence annotations from UniProt (31 features) and post-translational modification annotations from PhosphoSitePlus[1] database (six features) in **Fig. 4a** for missense variants, and in **Supplementary Fig. 9** for synonymous, nonsense, and frameshift variants.

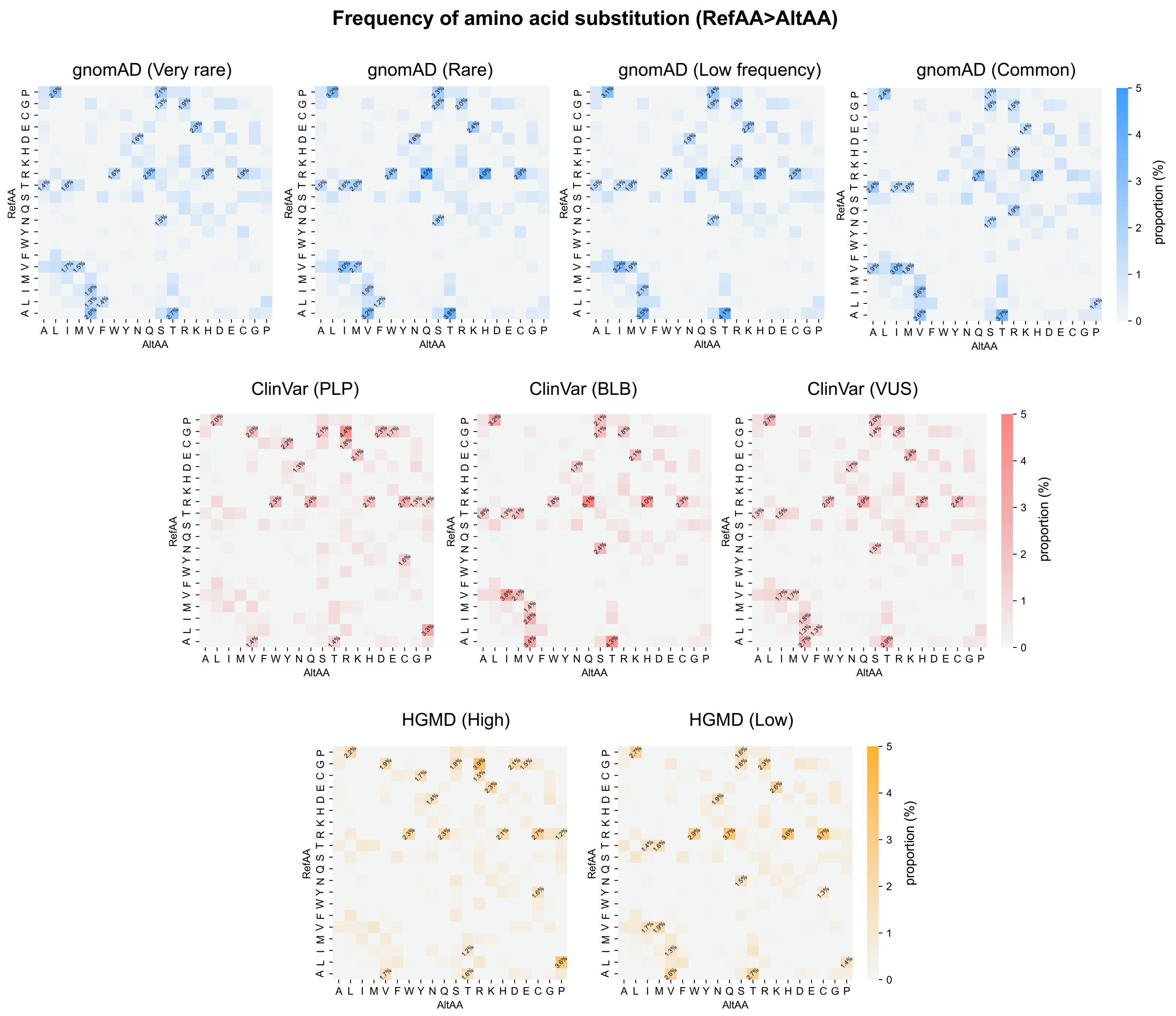


**Supplemental Fig. 8 | Amino Acid Change Frequencies in Missense Mutations:** This figure illustrates the frequency of amino acid changes resulting from missense mutations across nine variant datasets. Each missense variant is categorized by the specific amino acid substitution (e.g., Ala;A > Leu;L), encompassing 400 possible substitutions (20x20). The heatmap visually represents the proportion of each substitution, with the top 20 most populated substitutions quantified in each heatmap. We observe that gnomAD variants and ClinVar BLB variants exhibit high populations at specific amino acid substitutions, such as those occurring between aliphatic residues (V > A, I, I, or M), R>Q, A>T, or P>L. In contrast, pathogenic variant datasets (ClinVar PLP and HGMD High) demonstrate higher frequencies of substitutions involving special amino acids like G, C, and P. This observation suggests a discernible relationship between the physicochemical properties of reference and alternate amino acids in different variant datasets.

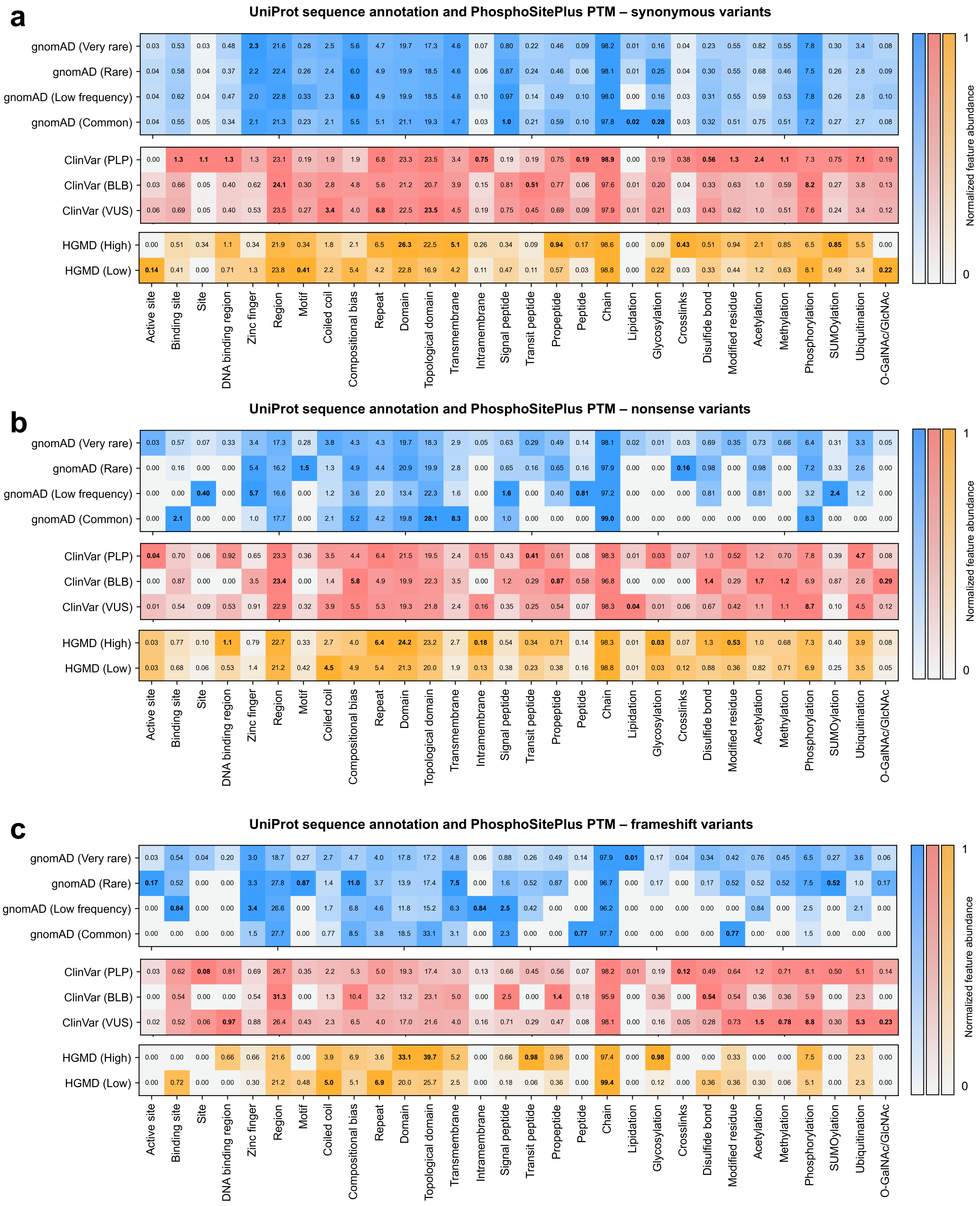


**Supplemental Fig. 9| Abundance of UniProtKB sequence annotations and PTM sites between nine variant datasets for synonymous, nonsense, and frameshift variants.** The figure replicates the calculations of feature abundance in missense variants shown in **Fig. 4a** for **(a)** synonymous, **(b)** nonsense, and **(d)** frameshift variants. The features abundance calculation method is described in **Methods** and **Supplementary Fig.** 7.

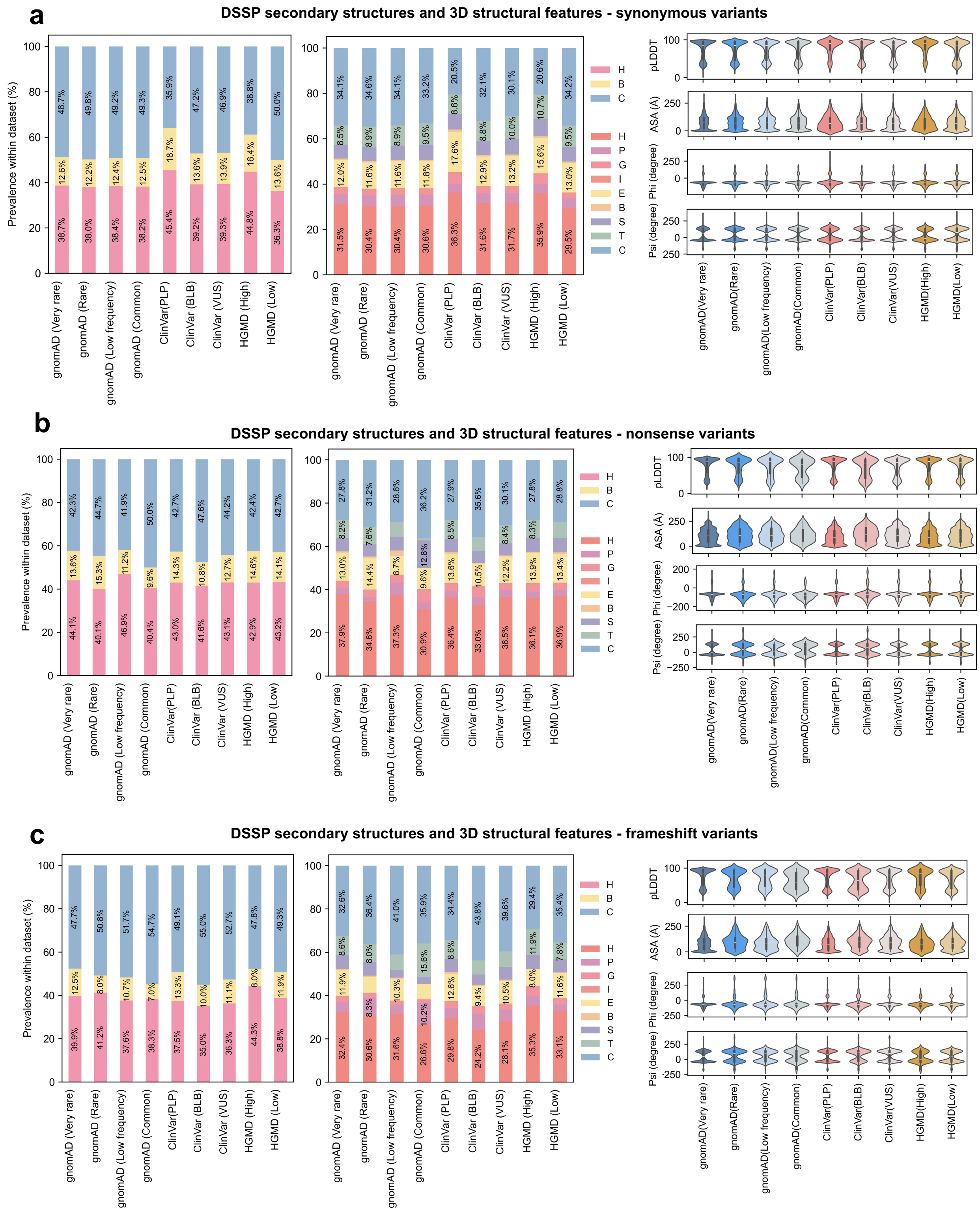


**Supplemental Fig. 10| Distribution of structural features across nine different sets of synonymous, nonsense, and frameshift variants.** The figure replicates the calculations from **Fig. 4b-c** for **(a)** synonymous, **(b)** nonsense, and **(d)** frameshift variants.

**Genomics 2 Protein Portal infrastructure**

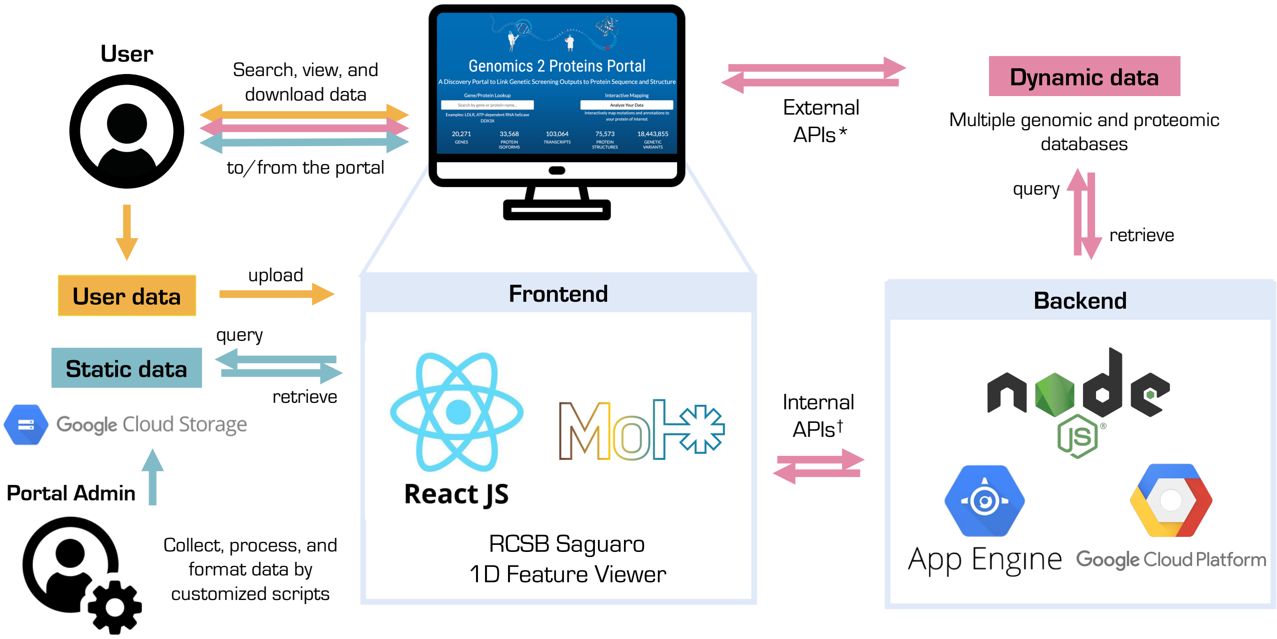


**† Internal APIs:**
List of genes: /api/genes/options
Get gene-wise meta data: /api/gene/:geneName
Get gene-family information: /api/geneFamily/:familyId
G2P3D API: /api/gene/:geneName/protein/:uniprotId/gene-transcript-map

***External APIs:**
Get protein sequence per uniprot id: https://www.ebi.ac.uk/proteins/api/proteins/:uniprotId
Get protein sequence per gene name: https://rest.uniprot.org/uniprotkb/search?format=fasta&query=:geneName
Get PDB structures: https://www.ebi.ac.uk/pdbe/graph-api/uniprot/unipdb/:uniprotId
Get AlphaFold structure: https://alphafold.ebi.ac.uk/api/prediction/:uniprotId
Get sequence annotations from UniProt: https://rest.uniprot.org/uniprotkb/:uniprotId.json
Get structure coverage for protein chains: https://www.ebi.ac.uk/pdbe/api/pdb/entry/polymer_coverage/:pdbId/chain/:chainId

**Static data:** variants from gnomAD, ClinVar, HGMD, protein features from PhosphoSitePlus, MAVEdb data, pre-computed structures features (secondary structures, accessible surface area, dihedral angles, etc.) using DSSP based on ALphaFold structures.

**Dynamic data:** protein sequences from UniProtKB, protein features from UniProtKB, structural features from SIFTS/PDBe, structures from PDB and AlphaFold

**Supplemental Fig. 11| The Google cloud infrastructure of the G2P portal.** This figure illustrates the web implementation of the portal. The frontend is implemented in React.js and includes RCSB Saguaro 1D Feature Viewer[2] and Mol* as protein sequence and structure viewer[3], respectively. The backend is implemented in Node.js and uses the Google app engine. Users can query, upload, and retrieve data from the portal, and the flow of user-uploaded, static, and dynamic data is shown with arrows in different colors (user-uploaded data in *orange*, static data in *cyan*, and dynamic data in *pink*). All static data are stored in Google cloud storage. All user-uploaded data remain on the users' browser, securing the confidentiality of users' data.

### Genomics 2 Protein Portal Sitemap

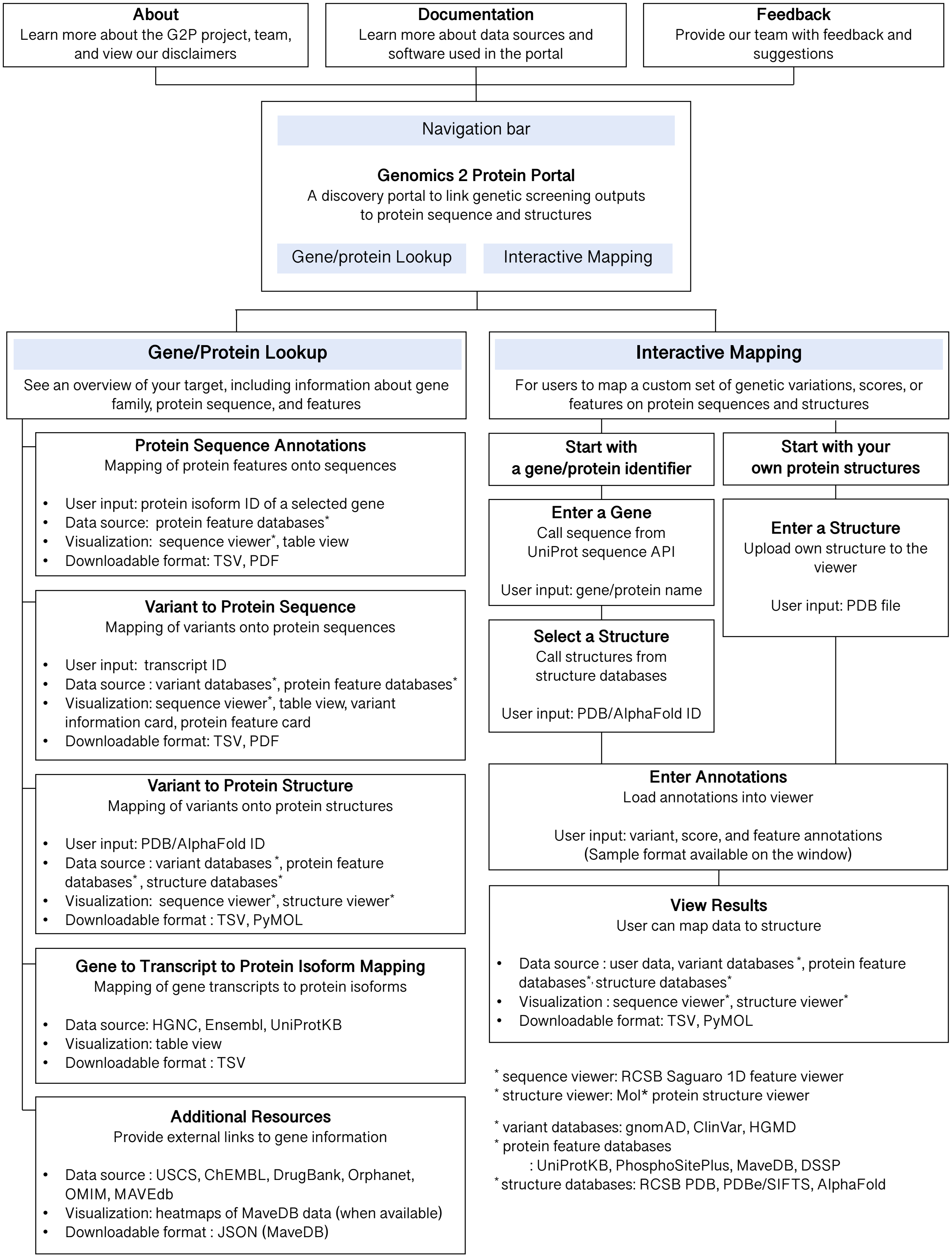

**Supplemental Fig. 12| A sitemap of the G2P portal.** From the home page, users can access the *About*, Documentation (*Doc*), and *Feedback* pages, available on the navigation bar at the top of the portal. There are two main modules in the portal: (1) Gene/Protein Lookup, accessible via searching by a human gene or protein name; (2) Interactive Mapping, accessible via secure Google sign-in upon clicking on the button displayed on the home page. The Gene/Protein lookup module has five submodules for *protein sequence annotation*, *variant mapping to protein sequence*, *variant mapping to protein structure*, *gene to transcript to protein isoform mapping*, and links to *additional resources*. The Interactive Mapping module has two submodules, for allowing users to start with any human gene or a protein structure to map user-uploaded data onto the target protein's sequence and structure. The user input, data sources, visualization methods, and downloadable data formats within each submodule are listed in the figure.

### Variant information and protein feature card

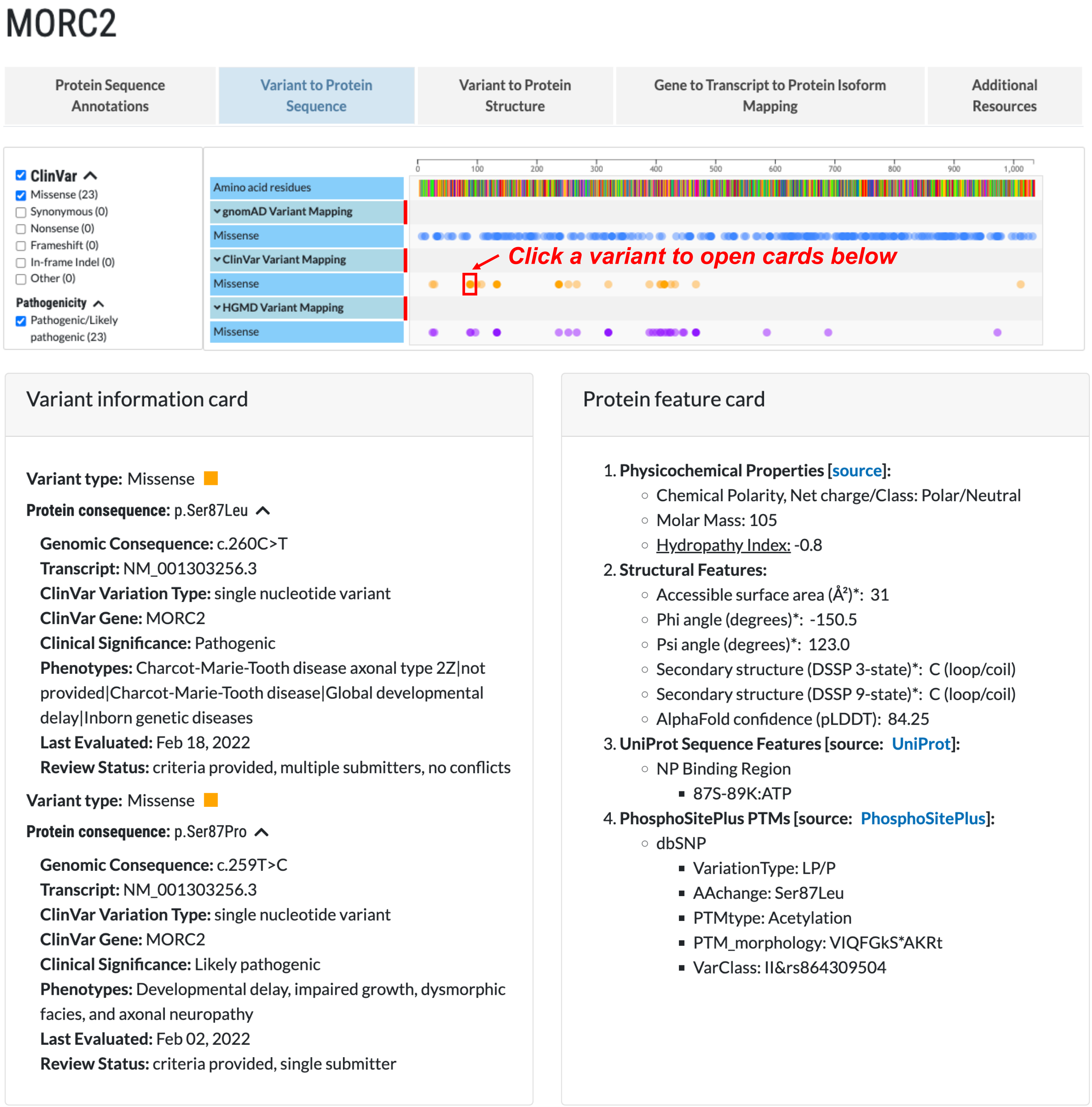


**Supplemental Fig. 13| Variant and protein feature card**. Upon selecting a variant on the sequence viewer (**Fig. 5a**), the variant information and protein features card will pop up, featuring detailed information about the selected genetic variant and a list of protein features on the variant position selected (see **Methods**). In the example shown here, *MORC2* ClinVar missense variant at Ser87 was selected, to view the cards revealing details for variants Ser87Leu and Ser87Pro, along with structural and functional annotations on Ser87.

### Mutagenesis output viewer

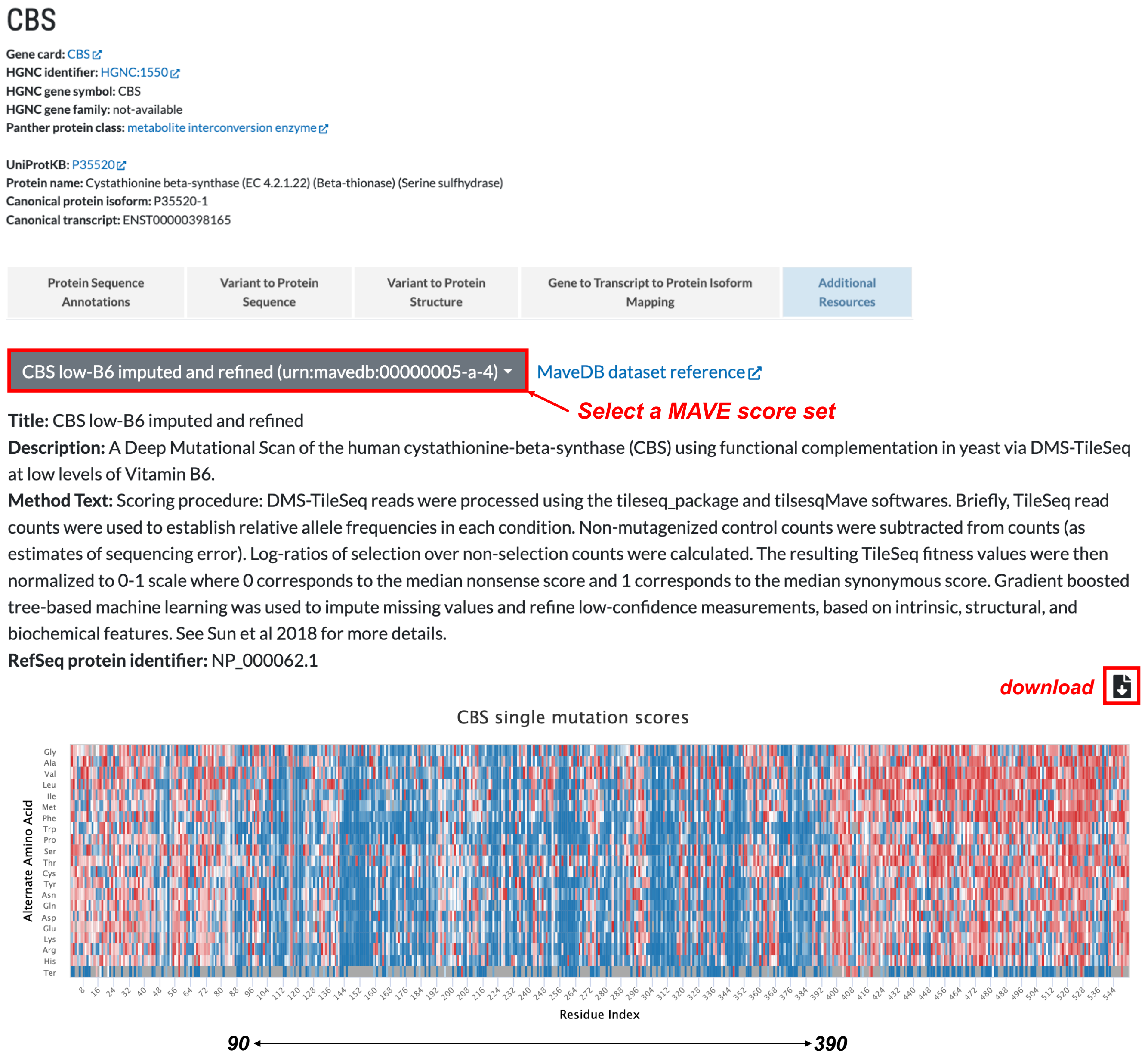


**Supplemental Fig. 14| Mutagenesis output viewer.** When a gene has available data in MaveDB[4], a brief description of a methodology and the assay readouts are shown in a heatmap. In the example illustrated in this figure (*CBS* MAVE readouts via DMS-TileSeq at low levels of Vitamin B6), scores show a clear distinction between residues 90-390 (low scores in blue) and residues at the N and C-terminus (high scores in red). By hovering over the heatmap, users can view the readouts from the assay and can download the entire score set by clicking the download icon.

### Supplemental Tables

**Supplemental Table 1| A list of genes in G2P3D API**. HGNC symbol, HGNC ID, locus group, and corresponding UniProtKB accession are given for 20,292 genes.

**Supplemental Table 2| HGNC gene family.** Count of genes for 834 HGNC gene families.

**Supplemental Table 3| PANTHER protein class.** Count of genes for 24 PANTHER protein classes

**Supplemental Table 4| Number of available PDB structures.** Count of PDB structures of 7,973 genes.

**Supplemental Table 5| A list of genes available from MaveDB.** 40 genes available in MaveDB and their scores are given.

**Supplemental Table 6| Annotations for interactive mapping case study.**
